# Protecting host mitochondria with vitamin B3 preserves bioenergetics and thermogenesis during antibiotic treatment

**DOI:** 10.64898/2026.11.08.765029

**Authors:** Tilman Tietz, Mügen Terzioglu, Laura Vesala, Tea Tuomela, Kabir Mahmud, Topi Paappanen, Pihla Lindroos, Eve Nyman, Kari Moisio, Juhana Juutila, Karita Karppinen, Kukka Lehto, Kirsi-Maarit Lehto, Susanna Valanne, Ville Hietakangas, Tiina S. Salminen

**Author notes:** Corresponding author: Tiina S. Salminen. shared authorship (Mügen Terzioglu, Laura Vesala).

## Abstract

Antibiotics that target bacterial translation can also impair mitochondrial function because mitochondria retain bacterial-like ribosomes. As antibiotic resistance continues to increase and the use of several broad-spectrum antibiotics has been limited by their mitochondrial toxicity, there is a growing need to understand their off-target effects in host physiology, and to identify strategies that can mitigate antibiotic-induced mitochondrial dysfunction. We demonstrate that antibiotic chloramphenicol (CA) induces a dose-dependent reduction in mitochondrial respiration and mitochondrial thermogenesis in *Drosophila* S2 cells, human HEK293T cells and in developing *Drosophila* larvae. *In vivo*, CA exposure triggers female-biased aberrant innate immune activation of hosts carrying naturally occurring mitochondrial rRNA variants. Metabolomic profiling revealed alterations in vitamin B_3_ metabolism in *Drosophila* during and following CA exposure. Among the vitamin B_3_ vitamers examined, nicotinamide (NAM) exhibited the strongest protective effect, mitigating the CA-induced impairments in mitochondrial respiration and thermogenesis, as well as aberrant immune activation. We identify mitochondrial thermogenesis as a previously unrecognized target of antibiotic-induced mitochondrial perturbation and a sensitive marker of mitochondrial stress. Furthermore, we demonstrate that co-supplementation of CA with NAM mitigates antibiotic-induced mitochondrial toxicity through evolutionarily conserved mitochondrial pathways. Finally, we establish *Drosophila* as a powerful model for investigating antibiotic-induced mitochondrial dysfunction and for developing therapeutic strategies to protect host mitochondria during antibiotic treatment.

## Introduction

The global rise in antibiotic resistance, driven by the widespread overuse and misuse of antibiotics in healthcare and agriculture, underscores the need to investigate the off-target effects of potent antibiotics that were previously abandoned or overlooked because of their adverse effects on host physiology. Because mitochondria, the ATP- and heat-producing organelles of our cells, are descendants of ancient bacteria^1,2^, they retain bacterial features that render them susceptible to antimicrobial drugs targeting pathogenic bacteria. Consequently, antibiotics can exert unintended off-target effects on mitochondrial function^3,4^, particularly during prolonger or high-dose treatment. Such mitochondrial perturbations may lead to severe cellular stress, underscoring the need for novel strategies to prevent or mitigate antibiotic-induced mitochondrial dysfunction. Moreover, mitochondrial perturbations are increasingly recognized as key contributors to a broad spectrum of diseases and syndromes, including metabolic disorders, neurodegenerative diseases, cancer, irritable bowel syndrome and many more^5–8^.

Mitochondria contain their own small, circular, maternally inherited genome (mtDNA). About 60% of approved antibiotics combat pathogenic bacteria by targeting ribosomes^9^ and their adverse effects can involve mitochondrial ribosomes (mitoribosomes), which are responsible for the translation of the 13 mtDNA-encoded peptides required for the assembly of oxidative phosphorylation (OXPHOS) complexes. Mitoribosomes retain a conserved rRNA core comprising the mitochondrial large and small ribosomal subunits (16S mt-rRNA and 12S mt-rRNA, respectively), as well as protein domains responsible for codon decoding and peptide bond formation^10,11^. These conserved functional elements are homologous to those of bacterial ribosomes and render mitoribosomes susceptible to antibiotics that target bacterial translation machinery. The broad-spectrum antibiotic chloramphenicol (CA), originally isolated from actinobacterium *Streptomyces venezuelae*^12^, and effective against many Gram-positive and Gram-negative bacteria^13–15^, was the first antibiotic produced entirely by chemical synthesis in 1949 at commercial scale, facilitating its rapid global availability^16^. It was initially used for serious infections such as typhoid fever, typhus and meningitis and became widely used due to its effectiveness. However, its use declined soon after the discovery of severe side effects in 1950’s, such as bone marrow toxicity^17,18^. CA binds to the peptidyl transferase centers (PTC) of the bacterial 50S ribosomal subunit^19^, but also inhibits mitochondrial translation by binding to the large subunit of mitochondrial ribosomes, leading to ribosome stalling and aberrant translation of mtDNA encoded proteins, decrease in the synthesis of electron transport chain (ETC) subunits, defective OXPHOS, depletion of ATP and metabolic dysfunction^15,20,21^. Thus, despite its efficacy against bacteria and favorable pharmacokinetic properties, CA is not recommended for the treatment of bacterial infections because of its severe adverse effects in the host, including hematopoietic disorders^22–24^.

Epistatic interactions have been reported between antibiotics, including CA, that impair pathogen protein translation, and the host mtDNA variants^25,26^. Recognition of these interactions would be important for minimizing the risk of iatrogenic harm, particularly given the widespread use of such antibiotics. Current understanding of the prevalence, mechanisms and clinical consequences of these interactions remain limited^27^, especially with respect to their effects on immune responses and overall host immunocompetence, as both mitoribosome-affecting antibiotics^28^, and certain mtDNA variants^29^ have independently been shown to promote immune activation. However, it can be challenging to investigate the systemic dose-dependent effects of antibiotics on the host physiology, as well as the contribution of mtDNA variation in experimental settings where the nuclear genetic background must be controlled.

We leveraged both *in vitro* (fruit fly *Drosophila melanogaster* and human cell lines) and *in vivo* (fruit fly) models to characterize the dose-dependent effects of CA on mitochondrial respiration, heat production and metabolic profiles to identify potential therapeutic targets for alleviating the CA caused mitochondrial toxicity. *Drosophila* exhibits a high degree of conservation in mitochondrial genome organization and function compared with vertebrates^30–32^, making it a well-suited model organism for this study. *Drosophila* has also proven advantageous for investigating the host-dependent developmental toxicity of various antibiotic compounds^33–35^. Moreover, *Drosophila* is a relevant model for hematopoietic research^36^ and immunometabolism^37^ due to wide conservation of molecular pathways and mechanisms governing the organismal immune responses^38,39^, and have been used in studies focusing on mitochondrial function in immune responses^29,40,41^.

We show that the effects of CA exposure are conserved between fruit fly and human, and are profoundly shaped by host genetic background, causing aberrant immune activation in genetically susceptible hosts. Importantly, we establish a therapeutic strategy capable of mitigating the detrimental effects of CA on mitochondrial function and organismal physiology.

## Materials and Methods

### *In vitro* respirometry

Concentration dependent effects of chloramphenicol (CA) on mitochondrial respiration were measured from Schneider 2 (S2) cells, originally derived from *D. melanogaster* Oregon R embryos^42^, and shown to maintain a hemocyte-like transcription profile^43^. S2 cells were cultured at 25°C in Schneider’s medium (Sigma S9895-1L) supplemented with 10% heat-inactivated FBS (Sigma F9665-500ML), and Pen/Strep (100 U/ml penicillin and 100 µg/ml streptomycin; Sigma P4333-100ML). Pen/Strep is routinely used in cell culture and it has no effect on cellular growth or mitochondrial respiration^44^. The culture was routinely tested for mycoplasma every 3 months and was diluted 1:6 with fresh medium in every 4-5 days. Respirometry was performed on a Hansatech Oxytherm on intact S2 cells with 1 x 10^7^ cells suspended in 1 ml 1x PBS at 25°C. For the dose dependent effect of CA (Sigma-Aldrich C0378), S2 cells were treated with 10, 20, 50, 75 and 100 μg/ml CA in 1 x PBS for 24 hours prior to oxygen consumption measurements, beside controls which were given the same volume of 1 x PBS. Per replicate oxygen consumption rates (nmol/ml/min) were measured on several occasions: upon sealing the well (basal respiration), after the addition of 2.5 µM oligomycin A (complex V inhibitor; leak respiration), followed by a titration of FCCP (3 x 1 µM; spare respiratory capacity) and once after the subsequent addition of 1 µM antimycin a (complex III inhibitor; non-mitochondrial respiration). For each sample mitochondrial respiration was isolated by subtraction of non-mitochondrial respiration and oxygen consumption rates were normalized against the corresponding control to yield the final relative mitochondrial respiration rate (Δ Oxygen consumption rate), analyzed with Oxygraph Plus software (Version 1.01).

Concentration dependent effect of CA was also performed on human embryonic kidney (HEK293T) cell lines ^45,46^. HEK293T cells were grown in DMEM (Sigma-Aldrich, D6546) supplemented with 10% heat-inactivated FBS, 100 U/mL of penicillin and streptomycin, and 1% L-glutamine in 10 cm dishes. The cell culture media was routinely tested for mycoplasma every 3 months and stock culture maintained by trypsinization twice a week. Respirometry was performed on a Hansatech Oxytherm with 5 x 10^6^ live cells suspended in 1 ml 1x PBS at 37°C. For the dose dependent effect of CA, HEK293T cells were trypsinized and transferred to a new dish. Cells were acclimatized for 24 hours and treated with 20 μg/ml and 100 μg/ml CA for 24 hours prior to oxygen consumption measurements alongside with controls. For the rescue experiments, HEK293T cells were co-supplemented with 100 μg/ml CA and 10, 20, 40, 50, 60 or 100 mM of nicotinamide (NAM) (Sigma-Aldrich N3376) or 100, 200 or 1000 μM niacin (Sigma-Aldrich N4126). After 24 hours of incubation cells are trypsinized, counted (NanoEnTek, Eve automatic cell counter) and 5 x 10^6^ live cells were pelleted by centrifugation at 250 g x 3 min, washed with prewarmed 1 x PBS and subjected to oxygen consumption measurements. Per replicate oxygen consumption rates (nmol/ml/min) were measured with five to six biological replicates following the same protocol as described above for S2 cells.

### *In vivo* respirometry

Respirometry was performed on *D. melanogaster* whole-larvae homogenates prepared from pooled 3^rd^ instar larvae (15 to 20 individuals per sample) reared on control food, or that supplemented with 0.2, 2 or 3 mg/ml CA or co-supplemented with 300 µg/ml NAM. Larvae were collected and rinsed thoroughly in water, patted dry and weighed. Larvae were Dounce-homogenized with 5 strokes in 500 µl homogenization buffer (250 mM sucrose, 2 mM EGTA, 5 mM Tris/Hcl, pH 7.4). Residual tissue was removed through a drop-in cell strainer (pluriSelect pluriStrainer Mini 100 µm) by centrifugation at 300 rcf for 3 min at 4°C. To initiate respirometry, 300 µl of the homogenate supernatant were added to the respirometry (Hansatech Oxytherm) well primed with 700 µl respiration buffer containing 225 mM sucrose, 75 mM mannitol, 10 mM Tris (pH 7.4), 10 mM KCl, 10 mM KH_2_PO_4_, 5 mM MgCl_2_, supplemented with 10 mg/ml BSA. Stabilized oxygen consumption rates were recorded by sealing the well after addition of the sample (basal respiration), followed by addition of substrates (L-proline, pyruvate, glutamate (all 2.5 mM) and ADP (0.5 mM) (“substrate-induced respiration”); addition of oligomycin (2.5 µM for ORT 237, 5 µM for ORT) (leak respiration); titration of FCCP (0.1, 0.2 and 0.3 µM; spare respiratory capacity or maximum respiration) and addition of antimycin a (1 µM, non-mitochondrial respiration). Oxygen consumption (nmol/ml/min) was normalized against sample biomass (nmol/ml/min/mg) or protein content by initially freezing 200 µl of the homogenate supernatant for protein quantification using the BCA protein assay kit (Merk Millipore). Mitochondrial respiration was calculated by subtracting non-mitochondrial respiration from “basal” and “maximum” respiration (after FCCP addition). Mitochondrial respiration was then normalized against the corresponding control to yield the final relative mitochondrial respiration rate (Δ Oxygen consumption rate).

### *In vitro* mitochondrial thermogenesis

Concentration dependent effect of CA on mitochondrial thermogenesis was measured with thermo-sensitive small chemical probe, Mito-Thermo-Yellow (MTY)^47^. Cells were prepared the same way as in respirometry: after treating cells with 20 μg/ml and 100 μg/ml CA for 24 hours, 1 x 10^7^ S2 cells and 5 x 10^6^ HEK293T cells were resuspended in 10 ml pre-warmed corresponding cell culture media for each cell type, supplemented with 100 nM MTY. Cells were incubated for 15 min at 25°C (S2 cells) or 37°C (HEK293T cells) and pelleted at 250g x 3 min. Cells were washed with pre-warmed 10 ml 1 x PBS, centrifuged at 250g x 3 min to remove the unbound chemical probe. PBS was removed and pelleted cells were kept about 10 minutes at 25°C/37°C incubator to deplete oxygen. Cells were then resuspended in 2 ml pre-warmed PBS, transferred into a quartz cuvette with continuous stirring and thermometric follow-up was performed on spectrofluorometer (PTI, Quanta Master TM, Horiba) with time-based program where signal is excited at 542 nm and constant detection of signal/sec are recorded at 562 nm emission wavelength. Temperature of quartz cuvette is controlled with both Peltier jacket that holds the cuvette and circulating water from temperature-controlled water bath (Julabo, Corio CD-200F). Mitochondrial thermogenic function of control cells or in response to the treatments (CA, CA+NAM) was measured via OXPHOS complex I (cI) inhibition with rotenone, data evaluation and temperature calculations done as previously described^48^ and reported here as a change in the temperature (-Δ).

### Fly genotypes

Two commonly used *D. melanogaster* lines *w^11^*^18^ (VDRC #60000) and *Oregon R* (designated *Oregon R Tampere*, ORT, owing to long-term maintenance at Tampere University) were used in most of the *in vivo* studies as indicated in the results. Cytoplasmic hybrid (cybrid) lines were generated by backcrossing the virgin female progeny of the mtDNA (mitotype) donor line with the males of the nuclear genome (nucleotype) donor lines^40^. Six mtDNA donor lines (**Table 1**) were selected from the *Drosophila* Genetic Reference Panel (DGRP)^49^ based on reported mitoribosomal mtDNA mutations that these lines possess^50^. To ensure near-complete nucleotype replacement the backcrossing regimen was maintained for at least 15 generations, before the cybrid lines were maintained independently.

**Table 1.** Mitochondrial genome (mtDNA) variants that contain variation in their mitochondrial ribosomal RNA genes (mt:rRNA) were selected from the Drosophila Genetic Reference Panel (DGRP). mtDNA naming is based on the DGRP mtDNA donor line identifier number. Studied mtDNA variants exist in their native nuclear background and in Oregon RT (ORT) and *w^1118^* nuclear backgrounds in the cytoplasmic hybrid (cybrid) lines. *mt:lrRNA* = *mitochondrial large ribosomal subunit*, *16S*; *mt:srRNA* = *mitochondrial small ribosomal subunit*, *12S*.

| **mtDNA** | **mt:rRNA variation** | **Nuclear genotype** |
|---|---|---|
| mt208 | *mt:lrRNA*: 12834_C/T and 13217_A/G | Native, ORT, *w*<sup>*1118*</sup> |
| mt237 | *mt:srRNA*: 14620_T/C | ORT, *w*<sup>*1118*</sup> |
| mt321 | *mt:lrRNA*: 13217_A/G | Native, ORT, *w*<sup>*1118*</sup> |
| mt377 | *mt:lrRNA*: 13562_A/G | Native, ORT, *w*<sup>*1118*</sup> |
| mt427 | *mt:lrRNA*: 12898_G/A | Native, ORT, *w*<sup>*1118*</sup> |
| mt765 | *mt:srRNA*: 14666_T/TTA | Native, ORT, *w*<sup>*1118*</sup> |
| mtORT | *mt:srRNA*: 238insTAA | ORT |
| *mtw*<sup>*1118*</sup> | None described | ORT, *w*<sup>*1118*</sup> |

For the OXPHOS gene knockdown via RNA interference (RNAi), we used the *in vivo* binary GAL4-UAS expression system^51^. RNAi construct expression was directed to hemocytes using a combination of two GAL4 driver lines, *Hml^Δ^-GAL4, EGFP* and *He-GAL4,GFPnls*^52^, referred to as *HH>*. An RNAi construct targeting the OXPHOS complex V gene *ATP synthase coupling factor 6* (*ATPsynCF6*; #107826/KK) and its corresponding background line *w^1118^* (#60100), were obtained from the Vienna *Drosophila* Resource Center (VDRC).

### Fly maintenance and preparation of supplemented food

Flies were reared and kept on potato-yeast control food or that supplemented with the indicated chemicals. The control food contained 36 g instant mash potato, 9g industrial agar, 45.5 ml white syrup, 14.5 g dry yeast, 8 g methyl-4-hydroxybenzoate (Nipagin) (Sigma-Aldrich H5501) and 5 g L-ascorbic acid (Sigma-Aldrich A7506) per a liter of water. Drug supplemented food was prepared by adding the finely powdered compounds (Chloramphenicol, Sigma-Aldrich C0378; Nicotinamide (NAM), Sigma-Aldrich N3376 or N-acetyl cysteine (NAC), Sigma-Aldrich A7250) directly to the cooling (<65°C) food to the specified final concentrations. Compound degradation was minimized by using the drug supplemented food within five days after preparation and avoiding exposure to direct sunlight. Flies and larvae were maintained at 25°C and 12:12h light:dark cycles for maintenance and experiments, except for the larval progeny of *ATPsynCF6* knockdown crosses, which were kept at 29°C.

### Egg-to-pupae development and pupariation success

The impact of different doses of CA on *Drosophila* larval development was screened at 25°C. 15-25 female and 8-10 male (<10 d) parental flies were mated per vial and after 24 hours the eggs were counted to control the effect of larval density on the development rate. Vials were scored regularly and per scoring event all newly formed pupae were counted. Pupariation success was calculated for each vial. Experiments were conducted as three independent blocks of three temporally staggered repeats.

### Melanotic nodule screen

3^rd^ instar larvae were collected, washed and dissected in a water droplet under the microscope. Larvae were scored as melanotic nodule-positive when at least one “black mass” of any size was found in the body cavity. In the initial screen (Fig. 3B-C), the sex of the larvae was not recorded. Where sex-specific data were collected, this information is indicated in the corresponding plots.

### Metabolome samples and analysis

Metabolome analysis was performed with adult flies and 3^rd^ instar larvae. For the fly samples, egg-to-larval stages were exposed to 2 mg/ml CA diet or reared on a control diet. Pupae were collected and transferred to control diet vials, without exposing them to CA after the eclosion. Two days old *w^1118^* and ORT males were collected in pools of five to obtain four biological replicates. For the larval samples, egg-to-3^rd^ instar larval stages were reared in 2 mg /ml CA supplemented food, food supplemented with CA and 300 µg/ml NAM or the control food. For each sample 15 3^rd^ instar larvae were checked for their melanotic nodule status, thoroughly washed and flash frozen. Larvae were scored as nodule-positive when at least one melanotic nodule was found in the body cavity.

The samples were analyzed on a Thermo Q Exactive Focus Quadrupole Orbitrap mass spectrometer coupled with a Thermo Dionex UltiMate 3000 HPLC system (Thermo Fisher Scientific). The HPLC was equipped with a hydrophilic ZIC-pHILIC column (150 × 2.1 mm, 5 μm) with a ZIC-pHILIC guard column (20 × 2.1 mm, 5 μm, Merck Sequant). Per sample, 5 µl were injected into the LC-MS instrument after quality controls in randomized order having every tenth sample as blank. A linear solvent gradient was applied in decreasing organic solvent (80–35%, 16 min) at 0.15 ml min–1 flow rate and 45 °C column oven temperature. Mobile phases were aqueous 200 mmol per liter ammonium bicarbonate solution (pH 9.3, adjusted with 25% ammonium hydroxide), 100% acetonitrile and 100% water. Ammonium bicarbonate solution was kept at 10% throughout the run, resulting in a steady 20 mmol per liter concentration.

Metabolites were analyzed using a mass spectrometer with a heated electrospray ionization source using polarity switching and the following settings: full-scan mode using range of 58 – 870 m/z and resolution of 70,000 at m/z of 200; spray voltages of 3,400 V for positive and 3,000 V for negative mode; sheath gas of 28 arbitrary units (AU) and auxiliary gas of 8 AU, vaporizer temperature of 280 °C and ion transfer tube temperature of 300 °C. The instrument was controlled using Xcalibur 4.7.102.25 software (Thermo Scientific). Metabolite peaks were confirmed using commercial standards (Sigma-Aldrich). Data quality was monitored throughout the run using an in-house quality control standard mix consisting of 76 commercial metabolite standards as well as an in-house quality control cell line extracted similarly to other samples. After final peak integration with TraceFinder 4.1 SP2 software (Thermo Scientific), peak area data was exported as Excel files. Analysis of peak area data was performed with the Metaboanalyst web service (Metaboanalyst.ca) using 10% IQR filtering, normalization by median, log10 transformation and auto-scaling (mean-centering and division by standard deviation for each variable). Hit metabolites were filtered by FC > 1.5 and p-value < 0.05.

### Flow Cytometry

We employed an Accuri C6 flow cytometer to quantify populations of circulating hemocytes in 3^rd^ instar larvae. Sample preparation workflows and event gating strategies were adapted to the larval genotype as described below. Larvae were collected and thoroughly rinsed in droplets of milliQ water to remove residual substrate. Larvae were then bled (hemolymph release) individually on a 12-well glass slide (Thermo Scientific) in 20 µl droplets of bleeding buffer (1 % bovine serum albumin (BSA; Sigma) in 1x Phosphate buffered saline (PBS); filtered through a 0.2 μm sterile syringe filter (VWR). This was done by carefully tearing the cuticle (dorsal side) along the posterior-anterior body axis with forceps (Fine Science Tools) and then carefully shaking the carcass to rinse the body cavity with bleeding buffer. After removal of the carcass sample droplets were carefully mixed and added to 80 µl bleeding buffer in 1.5 ml reaction tubes (yielding 100 µl of sample per larva). Per sample, 30 µl were run with the Accuri C6 at medium fluidics speed. Hemocytes were stained with propidium iodide (PI; Invitrogen F10797) (1 µl per 100 µl sample), and hemocyte quantification was performed as previously reported ^29^. Shortly, hemocytes were first separated from unspecific events based on a forward scatter area (FSC-A) vs. side scatter area (SSC-A) plot gate (**SFig. 1Ai**). Dead (PI-positive) cells were then excluded from the quantification (**SFig. 1Aii**). Live (PI-negative) cells were sub-gated into 2 morphological categories based on their positioning in forwards scatter area vs. forward scatter height plots (**SFig. 1Aiii**): The irregularly shaped activated hemocytes types such as lamellocytes deviate from the virtual correlation midline (45 ° angle) that characterizes round cells such as inactivated plasmatocytes and crystal cells ^53^. Hemocyte counts were back calculated to the full sample volume (100 µl (full sample) / 30 µl (sampled volume); rounded up). For each larva sex and nodule status were recorded.

**Figure 1.**
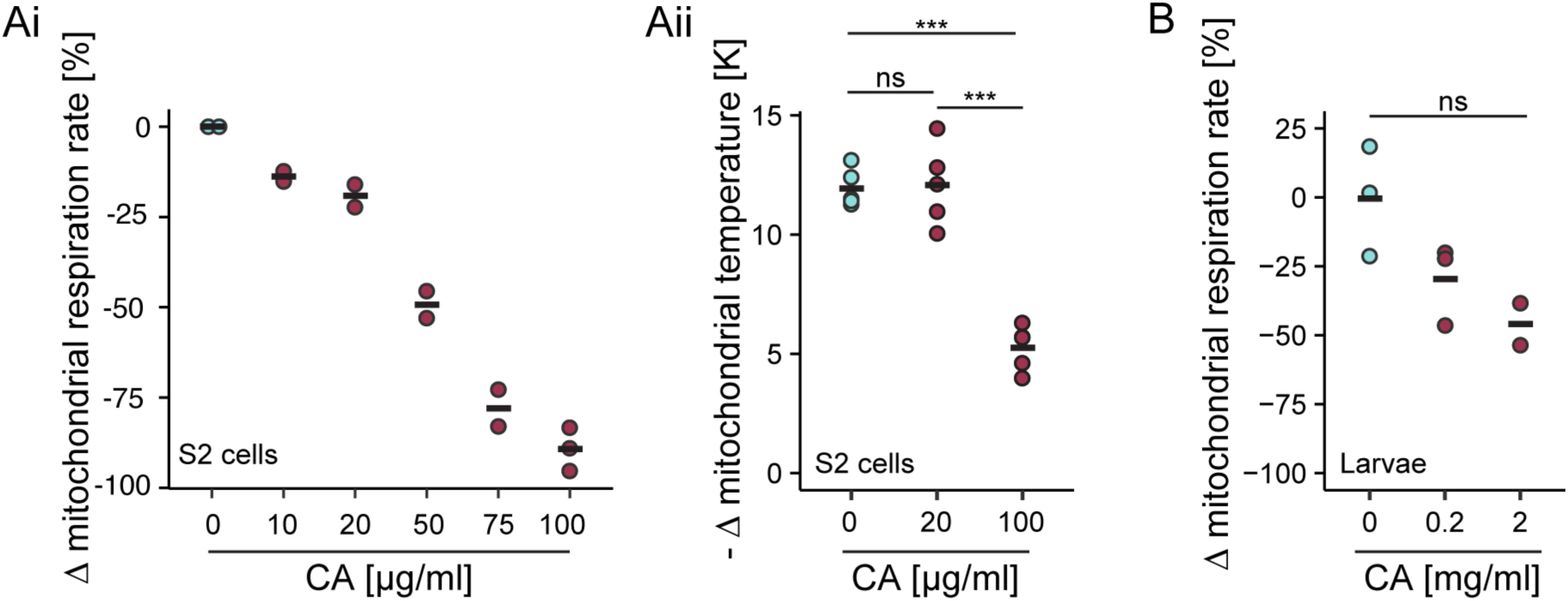
Chloramphenicol (CA) reduces mitochondrial respiration and decreases mitochondrial temperature. Ai) *Drosophila* S2 cells were exposed to CA for 24 hours, after which the oxygen consumption rate was measured and normalized to untreated control cells. Measurements were performed using 1 x 10^7^ cells per run in two to three replicates for each dose. **Aii)** Mitochondrial temperature was measured in control and CA-exposed cells using the thermosensitive fluorescent dye MitoThermo Yellow (MTY) (5 replicates per treatment). Low and high CA concentrations were selected based on the oxygen consumption. **B)** *D. melanogaster* larvae (ORT strain) were reared on control food or that supplemented with 0.2 or 2 mg/ml CA, and oxygen consumption rates were measured in 3^rd^ instar larvae. Normalization of the respiration data was performed against the sample biomass and change in respiration was calculated relative to control values. ns, not significant; * p< 0.05; ** p< 0.01; *** p< 0.001

### Quantifying larval crystal cells

Crystal cells were visualized by heating 11-17 3^rd^ instar larvae in 100 μl of 1 x PBS at 60°C for 10 minutes, which induces melanization of crystal cells and enables their direct counting ^29^. For imaging, larvae were mounted in a drop of glycerol on a glass slide and covered with a coverslip. Crystal cell numbers were determined from the whole larvae by using light microscope obtained images (example given in **Fig. 3Fi**) and the Fiji ImageJ (v1.52n) program.

### Immunofluorescent hemocyte imaging

Hemocyte imaging was carried out essentially as described previously^54,55^. We stained hemocyte samples with a lamellocyte-specific antibody to support flow cytometry data. 3^rd^ instar larvae were collected and thoroughly rinsed to remove residual substrate. Then, per well on a 12 well glass slide (Thermo Scientific), 2-3 larvae were successively bled in a 20 µl droplet of bleeding buffer (1 % bovine serum albumin (BSA; Sigma) in 1x Phosphate buffered saline (PBS); filtered through a 0.2 μm sterile syringe filter (VWR), by carefully tearing the cuticle (dorsal side) along the posterior-anterior body axis with forceps (Fine Science Tools) and carefully shaking the carcass to rinse the body cavity with bleeding buffer. Well slides were then incubated in a moisturized container for 45 min to allow for hemocyte attachment. Hemocytes were then fixed by the careful addition of 20 µl 4% paraformaldehyde (Invitrogen; v/v in 1x phosphate buffered saline) directly onto the bleeding buffer droplet. After 10 min of incubations cells were thoroughly washed with ice-cold PBS. The fixed cells were permeabilized then with 0.1% Triton X-100 for 5 min and again thoroughly washed with ice-cold PBS. Permeabilized cells were blocked with 3% BSA in PBS for 1 h at room temperature.

The affinity-purified monoclonal anti-L2 antibody (L2)^56^ was kindly provided by István Andó’s group (BRC, Szeged, Hungary). Anti-L2 specifically stains lamellocyte-lineage hemocytes. 20 µl of the L2 primary antibody was pipetted on hemocytes and incubated overnight at 4°C. The supernatant was then thoroughly washed off with ice-cold PBS. Secondary antibody staining was performed with 20 µl goat anti-mouse Alexa Fluor 488 secondary antibody (Life Technologies) in 3% BSA in PBS (1:500) for 1 h at room temperature in a light-proof container. The supernatant was then thoroughly washed off with ice-cold PBS. Cells were mounted with ProLong Diamond Antifade Mountant with DAPI (Life Technologies) and high-performance cover glasses (18 mm x 18 mm, Thickness no. 1.5; Zeiss) and cured overnight at room temperature in a light-proof container).

Immunofluorescent hemocytes were imaged using a Nikon A1R+ laser scanning confocal microscope with a Nikon Plan Apo VC 20x DIC N2, N.A. 0.75, WD 1.00 mm (Air) objective. Excitation was stimulated with built-in AOTF (Acousto-Optical Tunable Filter) lasers at 405 nm (DAPI) and 488 nm (Alexa Fluor 488). Fluorescence signals were filtered with 450/50 nm (DAPI), 482/35 nm (Alexa Fluor 488) emission filters.

The intensity of the Alexa Fluor 488 signal (L2 staining) was quantified from raw z-stack images using sum-intensity projections. For each image, the mean fluorescence intensity was measured and corrected by subtracting the mean background intensity. The background-corrected fluorescence intensity was subsequently divided by the number of DAPI-positive nuclei to account for differences in cell number between samples. Individual cell quantification was not feasible because lamellocytes frequently occurred in dense clusters. Image analysis and processing were performed using ImageJ2 (version 2.16.0/1.54p).

### Hemocyte RNA extraction

3^rd^ instar larvae were rinsed several times with filtered water and individually dissected in a drop of ice-cold 1 × PBS on a 12-well slide. Hemocytes from 50 larvae were pooled for each sample, and centrifuged at 2500 × g for 7 minutes at 4 °C. After centrifugation, excess PBS was carefully removed and the hemocytes were then snap-frozen on dry ice and stored at −80 °C. RNA was extracted using the Norgen Single Cell RNA Purification Kit (Norgen Biotek Corp., Ontario, Canada) with DNase treatment (RNase-Free DNase I Kit, Norgen). RNA concentration and purity were assessed using a NanoDrop ND-1000 spectrophotometer (Thermo Scientific).

### Reverse transcription quantitative real-time PCR (RT-qPCR)

RT-qPCR reactions were prepared with the SYBR Green real-time one-step qPCR kit (Thermo Fisher Scientific) using the following reaction mix: 5 µl iTaq Universal SYBR Green Reaction Mix (2×), 0.125 µl iScript reverse transcriptase, 0.3 µl each of 10 µM forward and reverse primers, 2.275 µl nuclease-free water, and 2 µl RNA (10 ng/µl). Reactions were run on a Bio-Rad CFX96 system with the following cycling protocol: reverse transcription at 50 °C for 10 minutes; polymerase activation and initial denaturation at 95 °C for 1 minute; followed by 36 cycles of 95 °C for 10 seconds and 60 °C for 15 seconds. A melt-curve analysis was performed at the end of the run in 0.5 °C increments from 65 °C to 95 °C, holding 2 seconds at each step. Experiments comprised three biological replicates with two technical replicates each. The expression ratios of target-to-reference gene (*Histone H3.3B*) in control and treatment groups were calculated using the dCT-method (2^^-dCT^). *ATPsynCF6* primer sequences were forward, 5’-CTGGACAAAGTGCGCGAGTA-3’ and reverse, 5’-CACGGTCCAGTTCGGTCTTC-3’ and *His3.3B* sequences were forward, 5’-CGCTCAGGATTTCAAGACCG-3’ and reverse 5’-GGATGTCCTTAGGCATGATTGT-3’.

### Transgenerational effects of CA

Parental flies were placed on either control or 0.2 mg/ml CA food to lay eggs, and the resulting progenies were collected as virgin females and young males and placed to feed on control food three-five days prior crossing. Crosses were done in four configurations, all on control food: I) females reared on control food crossed with males reared on control food, II) females reared on CA food crossed with males reared on control food, III) females reared on control food crossed with males reared on CA food and IV) females reared on CA food crossed with males reared on CA food. Progenies of these crosses were checked for melanotic nodule formation as described above.

### Antibacterial activity of CA, NAM and niacin alone or in combination

*Staphylococcus aureus* ATCC29213 and *Escherichia coli* K12 were grown overnight in Luria Bertani (LB) medium at 37°C and subcultured in fresh medium the next morning. Bacteria were grown to OD_600_ of 0.4-0.6, after which the bacterial suspensions were diluted 1:3 in fresh LB medium and spread on pre-warmed LB plates (200 µl of bacterial suspension/plate). After the suspension had completely dried, small filter paper discs were placed on top, to mark the spot where to insert each chemical. 50 µl drops of tested chemicals were carefully pipetted onto the filter paper discs and allowed to immerse and dry on the discs and the LB plate. The tested chemicals and their concentrations were as follows: Chloramphenicol (CA, Sigma-Aldrich C0378) 100 µg/µl, Nicotinamide (NAM, Sigma-Aldrich N3376) 40 mM and Niacin (Sigma-Aldrich N4126) 100 µM. Plates were incubated overnight at 37°C, allowing growth of the culture on the plate. Plates were photographed, and the areas showing clearance of bacteria around each chemical were estimated by using Fiji ImageJ (v1.52n) program measuring tools.

### HEK293T cell viability

The effect of CA on cell viability was determined with the 100 μg/ml CA dose and 17, 24, 30, 48, 72, and 96 hrs exposure intervals. HEK293T cells were counted and seeded as follows; for 17-, 24-, and 30-hour time points starting cell number is 700 000 and 40, 48, 72, and 96 hours starting cell number is 500 000. The CA exposure of the cells was started after 24 hours of seeding. For every time-point, exposed and control groups were prepared as triplicates on a 6-well plate. At the end of exposure time triplicate control and exposed groups were trypsinized and counted on Eve^TM^ Automated Cell Counter (NanoEntek). For every time-point cell viability and total number of cells were determined.

### Statistical analyses

The data on respiration (oxygen consumption) and mitochondrial temperature were analyzed using 2-tailed t-tests assuming equal variances in Microsoft excel. R studio^57^ was used for other data analyses and for plotting. For development time, counts were transformed into the approximate time to pupariation for each pupa, and data was handled by pairwise_survdiff from survival package^58,59^, calling a log-rank (G^\rho family) test with default Benjamini-Hochberg corrections. The pupariation success, melanotic nodule prevalence and cell survival was analyzed using a generalized linear model (GLM) with a binomial error distribution and logit link function. The response variable consisted of the number of eggs developed into pupae (success) and remaining number of eggs (failure), larvae with (nod^+^) and without (nod^-^) nodules or live and dead cells. Mitotype and CA treatment were included as fixed effects in the model. Mean fluorescence intensity was analyzed using non-parametric Kruskal-Wallis test followed by Dunn’s test for pairwise comparison. The number of hemocytes was analyzed using a GLM with a negative binomial error distribution to account for overdispersion in the count data. Mitotype and CA treatment were included as fixed effects. To compare estimated survival probabilities among mitotype-treatment combinations, estimated marginal means (EMMs) were calculated, and pairwise comparisons were performed with appropriate multiple-comparison adjustment. Lme4^60^, MASS^61^ and emmeans^62^ packages were utilized. In general, ggplot2^63^ was used for plotting, and Adobe Illustrator was used for figure editing. Respirometry and thermometry data values are in **Table S1**, data for pupariation rate and success, melanotic nodules, hemocyte counts and RT-qPCR in **Table S2** and filtered metabolite data in **Table S3.** Raw and normalized metabolite peak values are in **Table S4.**

## Results

### Chloramphenicol (CA) affects mitochondrial respiration and thermogenesis dose-dependently

It has been established that CA impairs metazoan mitochondrial protein translation^22,64^, and this impairment can lead to compromised mitochondrial respiration and reduced ATP synthesis in mammalian cell lines^20,65^. However, to our knowledge, the effects of CA on mitochondrial respiration in *in vivo* models, including *Drosophila,* and on the mitochondrial thermogenesis in general have not been investigated. During mitochondrial respiration, the electrons acquired from substrates are passed in ETC through multisubunit protein complexes while protons are pumped across the inner mitochondrial membrane via complexes I, III and IV. This electrochemical gradient produces ATP as well as heat, and physiologically active mitochondria with functional respiratory capacity have been shown to maintain 15°C higher temperature than the surrounding cell material^66^. This phenomenon was observed across multiple cell lines derived from both endothermic species (mouse and human) and a poikilothermic species, the fruit fly (*Drosophila melanogaster*)^48^.

To study whether CA affects *D. melanogaster* mitochondrial functions, we initially measured respiration and mitochondrial inner membrane temperature from CA exposed intact *Drosophila* S2 cells (**Fig. 1Ai-Aii**). A 24-hour exposure of S2 cells to CA at concentrations of 10, 20, 50, 75, and 100 µg/ml resulted in a dose-dependent reduction in mitochondrial respiration (**Fig. 1Ai**). We next used the mitochondrial heat production as an alternative readout of mitochondrial function upon CA exposure and selected lower and higher CA concentrations that produced a subtle or strong effect on mitochondrial respiration (**Fig. 1Ai**). Treating S2 cells with the low concentration (20 µg/ml) of CA for 24 hours caused 25% decrease in oxygen consumption (**Fig. 1Ai**) without altering the mitochondrial temperature (**Fig. 1Aii**). However, 100 µg/ml CA resulted in drastic (>80%) reduction in mitochondrial respiration (**Fig. 1Ai**) and significantly decreased the mitochondrial temperature when compared to the untreated S2 cells (**Fig. 1Aii**). To examine if similar dose-dependent effects on mitochondrial functions can be seen *in vivo*, we supplemented the *Drosophila* food with CA at concentrations of 0,2 and 2 mg/ml, reared the flies on control or CA supplemented food since the egg stage and measured respiration in pooled whole-larvae homogenates of the Oregon R (ORT) strain. Larval respiration showed a dose-dependent trend in response to CA exposure (**Fig. 1B**). These experiments were subsequently repeated in different experimental contexts, where CA induced significant mitochondrial respiratory defects (**Fig. 5A** and **Fig. S2**). Taken together, the *in vitro* and *in vivo* findings demonstrate that CA impairs mitochondrial function in *Drosophila*.

### Larval CA exposure leaves a lasting metabolic signature in unexposed adults

Since CA disrupts mitochondrial respiration and thermogenesis (**Fig. 1**), we next examined its phenotypic effects with two commonly used *D. melanogaster* lines, *w^1118^* and ORT. Supplementing the food with 1, 2 or 3 mg/ml CA did not delay the onset of pupariation (**Fig. 2Ai**), nor decreased the pupariation success (**Fig. 2Aii**), suggesting that the doses between 1-3 mg/ml CA do not cause developmental toxicity in *Drosophila*. In contrast, CA caused a moderate improvement in larval development and pupariation success in the ORT line (**Fig. 2Ai**). Overall, ORT larvae exhibited slower development and lower pupariation success than in the *w^1118^* line under control conditions and when exposed to CA.

**Figure 2.**
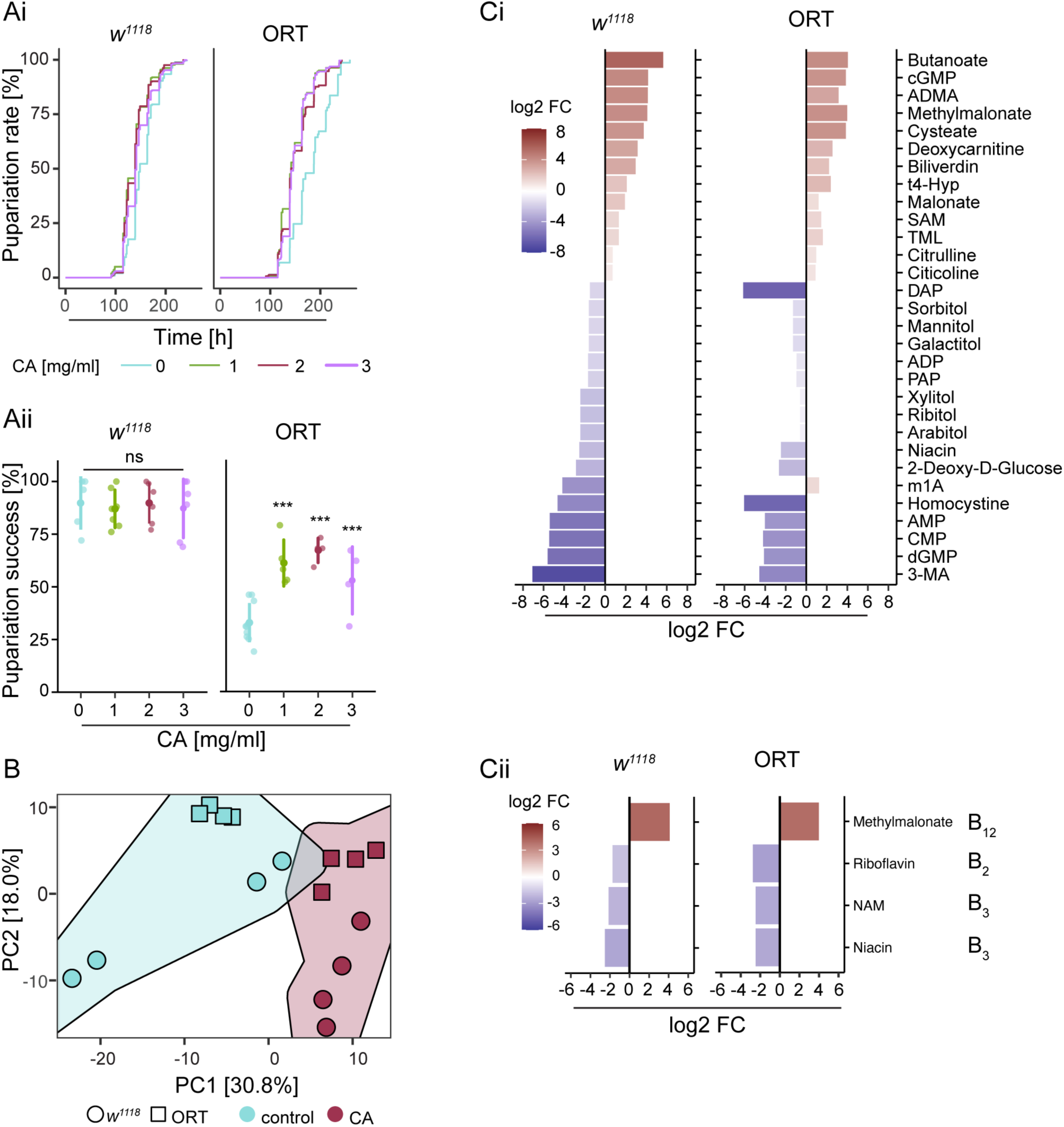
Larval CA exposure leaves a lasting metabolic signature in unexposed adults. Ai) Pupariation rate and **Aii)** pupariation success of CA exposed *w^1118^* and ORT larvae. **B)** PCA plot of the metabolome of unexposed adult male flies, reared from egg to larval stages on control food or that supplemented with 2 mg/ml CA (n = 4, pools of five flies). **Ci)** The 30 most significantly altered metabolites shared between *w^1118^* (p<0.0012) and ORT adults following larval CA exposure, ranked according to statistical significance in *w^1118^*. **Cii)** Relative abundances of significantly altered vitamin B and vitamin B-associated metabolites in *w^1118^*and ORT flies following larval CA treatment, relative to control-fed flies. ns, not significant; * p< 0.05; ** p< 0.01; *** p< 0.001

**Figure 3.**
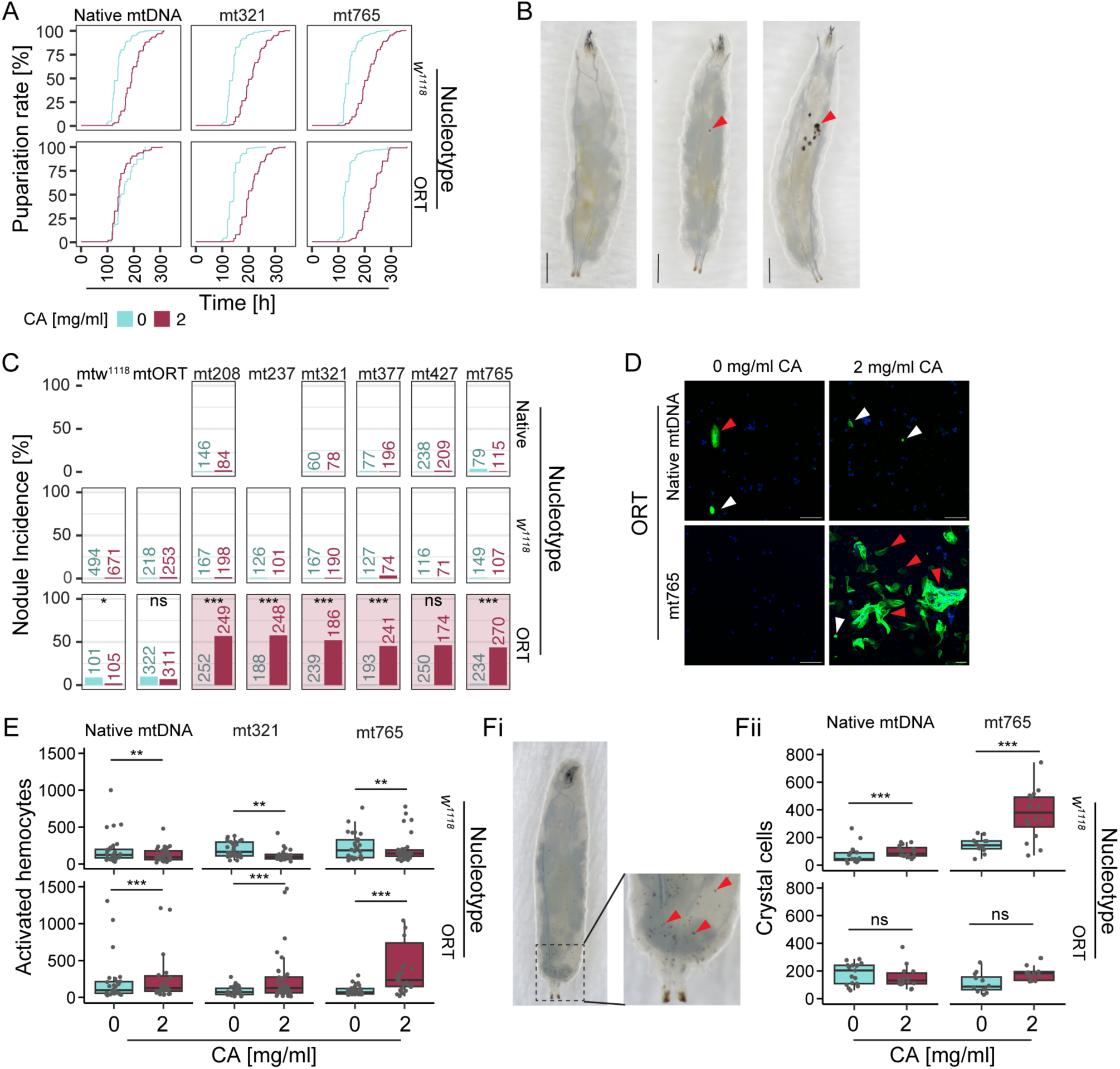
CA affects immune cell phenotype in a genotype-dependent manner. **A**) Pupariation rate of native ORT and *w^1118^* larvae and those of mt321 and mt765 cybrids reared at 2 mg/ml CA diet. **B)** 3^rd^ instar larvae with (left to right) no visible nodules, small nodule and multiple nodules. Nodules are indicated by red arrowheads. **C**) Percentage of nodule-bearing native and cybrid larvae per genotype x treatment cohort. Numbers of dissected larvae are above their corresponding plots. Statistical analyses were performed only on ORT cybrids due to the low melanotic nodule incidence in other nucleotypes. Panels with nodule incidences >10% are highlighted by red shading. **D)** Immunostaining of immune cells (hemocytes) collected from ORT and ORT-mt765 3^rd^ instar larvae reared on either a control diet (0 mg/ml CA), or a diet supplemented with 2 mg/ml CA. Images show fixed immunofluorescent hemocytes with DAPI stained cellular nuclei (blue) and L2 lamellocyte antibody (AlexaFluor488nm, green). Examples of L2-positive lamellocytes and lamellocyte aggregates are marked with red arrowheads, L2-positive hemocytes with non-lamellocyte morphology with white arrowheads. **E)** Number of immune activated lamellocyte-like cells from 3^rd^ instar larvae collected from control or 2mg/ml CA diet. 30 larvae were dissected per group. **Fi)** Representative light microscopy images of the crystal cells shown as black punctate (indicated by red arrowheads) and counted based on the whole larval images, **Fii)** and their quantification. ns, not significant; * p< 0.05; ** p< 0.01; *** p< 0.001.

Based on the respirometry and the phenotypic assays, we selected a CA concentration of 2 mg/ml for subsequent studies investigating the persistence of CA-induced effects on the metabolome of unexposed young adult *w^1118^*and ORT flies exposed to CA throughout larval development. These flies were not exposed to CA after the larval stages. Principal component analysis (PCA, **Fig. 2B**) of the metabolome showed stronger clustering based on the treatment (CA vs. control; PCA1; 30.8%) than based on the genotype (PCA2; 18%). Next, we focused on top 30 metabolites that were significantly altered in both *w^1118^* and ORT flies following larval CA exposure. These metabolites were ranked according to their statistical significance in *w^1118^* (**Fig. 2Ci**). Several metabolites remained elevated in both *w^1118^* and ORT flies, suggesting that CA induces long-lasting physiological and metabolic alterations that persist into adulthood after developmental exposure (**Fig. 2Ci**). The differentially abundant metabolites suggested perturbations in mitochondrial metabolism, including increased levels of methylmalonate^67^, a biomarker of oxidative stress, aging and mitochondrial dysfunction; malonate^68^, which is associated with impaired mitochondrial malonate-to-malonyl-CoA conversion, and the carnitine biosynthesis intermediates deoxycarnitine and N,N,N-trimethyllysine (TML)^69^. As carnitine biosynthesis pathway generates L-carnitine required for the transport of long-chain fatty acids into mitochondria for β-oxidation, these changes could suggest increased fatty acid transport to mitochondria. Metabolites that suggested oxidative stress included the simultaneous increase of the second messenger cyclic GMP (cGMP) downstream of nitric oxide (NO) signaling^70^, N,N-dimethylarginine (ADMA), an inhibitor of nitric oxide synthase^71^ and oxidative stress markers cysteate and biliverdin. Interestingly, simultaneous increase in the major methyl donor S-adenosylmethionine (SAM)^72^ and ADMA, released when methylated proteins are degraded, could indicate changes in methylation patterns. The metabolite data also indicated a purine and nucleotide depletion in CA-treated animals, with reduced levels of AMP, ADP, CMP, dGMP, 1-methyladenosine (m1A, down in *w^11^*^18^, up in ORT) and 3-methyladenine (3-MA) (**Fig. 2Ci**). These changes strongly indicate not only reduced nucleotide turnover, but also disturbance of cellular energy metabolism, which could indicate a low growth state. In addition, the concurrent decrease in sugar polyols (sorbitol, mannitol, galactitol, arabitol, ribitol and xylitol) may be consistent with reduced PPP activity, potentially reflecting lower NADPH availability for the reduction of sugars to polyols. The most upregulated shared metabolite altered in both genotypes was butanoate, a well-known microbial fermentation product. This together with downregulation of diaminopimelate (DAP), a cell wall component in most gram-, and some gram+ bacteria, could indicate altered host-microbe interactions and microbial metabolic output brought upon by CA feeding at larval stages.

When examining the 30 most significantly altered metabolites separately in the two backgrounds, ORT background showed additional indications of oxidative stress/altered redox homeostasis, whereas the *w^1118^* background exhibited signatures of lipid and methyl group metabolism (**Table S3**). Interestingly, *w^1118^* flies showed increased levels of 3-hydroxykynurenine (3-HK) and kynurenate, as well as xanthurenate, consistent with a shift of tryptophan metabolism toward the kynurenine pathway (**Table S3**).

B vitamins are essential cofactors facilitating cellular processes, including mitochondrial functions^73^, and altered levels of B vitamins and B-vitamin-derived metabolites are a common consequence of mitochondrial dysfunction. However, as the ability to synthesize B vitamins is widespread among commensal gut microbes, and CA is a broad-spectrum antibiotic capable of altering microbiota composition, CA treatment has the potential to change the levels of B-vitamin-related metabolites, in addition to its direct effects on mitochondrial functions. We found that several vitamin B-associated metabolites were affected by the CA-treatment also in *Drosophila* (**Fig. 2Cii**). Levels of two vitamin B_3_ vitamers, nicotinamide (NAM) and niacin, were significantly reduced in both *w^1118^* and ORT flies exposed to CA within the larval stages (**Fig. 2Cii**). Since B_3_ vitamers are nicotinamide adenine nucleotide (NAD^+^) precursors, and NAD^+^ is the primary electron carrier in cellular metabolism, reduced B_3_ levels may indicate increased demand for NAD^+^, which is common during various cellular stress conditions, including mitochondrial stress^74^. Another vitamin B with decreased levels was vitamin B_2,_ riboflavin, a precursor of the cofactors flavin mononucleotide (FMN) and flavin adenine dinucleotide (FAD), essential for oxidation-reduction reactions in the cell. Moreover, FAD, reduced in *w^1118^* after CA treatment (**Table S3**), is a cofactor of succinate dehydrogenase (OXPHOS complex II), multiple enzymes involved in fatty acid β-oxidation and the conversion of the amino acid tryptophan to niacin ^75^. Methylmalonate, a metabolite closely associated with vitamin B_12_ metabolism and a functional biomarker of vitamin B_12_ deficiency, was elevated in both *w^1118^* and ORT flies following larval-stage CA exposure (**Fig. 2Cii**). Among these metabolites, niacin and methylmalonate were also present in the top40 metabolite list (**Fig. 2Ci**).

Taken together, CA exposure did not cause overt developmental toxicity from the egg to pupal stage in our *in vivo* model. However, the metabolomic profiles of adults that had been exposed to CA only during the larval stage revealed persistent metabolomic alterations, indicating that CA exerts effects that carry over developmental stages and may have long-term physiological consequences.

### CA causes immune activation in specific mitonuclear combinations

Because CA treatment altered mitochondrial function both *in vitro* and *in vivo* (**Fig. 1**) and induced a metabolomic stress signature without causing overt developmental defects in flies, we next sought to investigate the effects of CA exposure across genetically diverse nuclear and mitochondrial genome backgrounds. We studied the potential additive effects of CA-induced and genetically originating mitoribosomal stress by exposing cytoplasmic hybrid (cybrid) lines carrying mitoribosomal mutations to CA. For this, we backcrossed six naturally occurring mtDNA variants obtained from the Drosophila Genetic Reference Panel (DGRP^76^) to *w^1118^* and ORT backgrounds, and gained 12 cybrid lines. Each of these mtDNA variants contain single nucleotide polymorphisms or insertions/deletions in their *12S* (*mt:srRNA; mitochondrial small ribosomal subunit*) or *16S* (*mt:lrRNA; mitochondrial large ribosomal subunit*) genes (**Table 1**)^76^. Inspecting the effect of 2 mg/ml CA supplemented diet on the pupariation rate of cybrid lines showed a prolonged onset of pupariation, apart from the native ORT line (**Fig. 3Ai**), showing that the CA treatment causes mild to moderate stress to the *w^1118^*and ORT cybrid flies.

*Drosophila* relies on cell-mediated and humoral innate immune responses^77^. In cell-mediated innate immunity, *Drosophila* immune cells, hemocytes, perform several essential functions in uninfected and infected animals, including phagocytosis of bacteria and apoptotic corpses (by plasmatocytes), melanin production and wound healing (by crystal cells), and melanin production together with encapsulation of foreign material (by lamellocytes at larval stages). In *D. melanogaster* larvae, hemocyte differentiation occurs on demand within a specialized organ called the lymph gland, and mature plasmatocytes can also transdifferentiate into other hemocyte types ^54,78–81^. We have previously reported that one of the outcomes of systemic or *Drosophila* hemocyte targeted mitochondrial perturbation is the aberrant activation of cell-mediated innate immunity and the emergence of melanized hemocyte aggregates called melanotic nodules^29,41,82^. Here, we screened the presence of melanotic nodules (examples shown in **Fig. 3B**) from the native DGRP lines and from the *w^1118^*and ORT cybrid lines reared on control or CA (2 mg/ml) supplemented food. Whereas we observed only modest incidence (<10%) of melanotic nodules in 3^rd^ instar larvae reared on control food, there was a significant and consistent increase in melanotic nodules in CA food reared ORT cybrid larvae, but not in the native ORT larvae (**Fig. 3C**).

Melanotic nodules are formed by an immune-activated cell type known as the lamellocyte, a specialized effector cell class distinguished from other immune cells by its larger size and characteristic discoidal shape^83^. Lamellocytes are rare or absent under baseline conditions in circulation, but emerge rapidly following immune insults such as wasp parasitation and cuticular injury^84,85^. To determine whether CA treatment triggers immune cell activation, larval hemocytes were examined by immunostaining and flow cytometry. We first focused on the nodule phenotype observed in the ORT cybrids and investigated the potential involvement of immune activation by imaging hemocytes from ORT and the nodule-prone cybrid line ORT-mt765 larvae. While ORT larvae reared on control food had occasional L2-positive hemocytes, they were reduced in CA-treated larvae. Notably, immune-activated hemocytes were consistently detected in CA-exposed ORT-mt765 larvae, frequently forming large aggregates (**Fig. 3D, Fig. S1B**). Next, we profiled the hemocytes of control and CA exposed native and *w^1118^* and ORT cybrid lines with flow cytometry. Also with this method, the CA treated nodule-prone ORT cybrid larvae exhibited elevated abundances of immune-activated lamellocyte-class cells (**Fig. 3E)**. However, the CA induced expansion was not observed in *w^1118^* derived cybrids, but instead the larvae reared on the control diet had elevated counts of activated hemocytes, which were rescued with the CA treatment (**Fig. 3E**). While CA exposure led to a decrease also in total hemocyte counts in *w^1118^,* in ORT they were increased, except of mt321 (**Fig. S1C**). Furthermore, CA exposure resulted in a slight increase in the proportion of dead hemocytes in *w^1118^* and ORT flies harboring their native mtDNA, while the opposite effect was observed in cybrids of both genetic backgrounds (**Fig. S1D**). Because the crystal cells cannot be recognized with our flow cytometry gating approach, we estimated the amounts of crystal cells from the whole larvae samples by quantifying the black punctate (example given in **Fig. 3Fi**) representing the ruptured crystal cells after heat treatment. Crystal cells were quantified from ORT-mt765 and *w^1118^*-mt765 cybrids and the ORT and *w^1118^* background lines reared with control and CA supplemented diet. In ORT and ORT-mt765 larvae the CA treatment did not affect the number of crystal cells, whereas in *w^1118^* and *w^1118^*-mt765 larvae the CA exposure increased the amount of crystal cells (**Fig. 3Fii**).

To summarize, exposure to CA causes distinct, but quite opposing immune cell phenotypes depending on the (cybrid lines’) genetic background. This is demonstrated by the formation of melanotic nodules, elevated amounts of immune activated hemocytes and unaltered levels of crystal cells in the ORT cybrids, whereas in *w^1118^* cybrids there were no melanotic nodules, amount of immune activated and total hemocytes were decreased and the amount of crystal cells was increased following the CA exposure.

### Melanotic nodule phenotype is female biased but is transmitted to the next generation via males

Because we observed a strong metabolic fingerprint in adult flies exposed to CA during the egg-to-larval stages, we next examined whether the CA-induced effects persist even longer. For this, we examined whether the nodule phenotype can be transmitted transgenerationally. To reduce possible residual CA levels after initial exposure, we decreased the concentration from 2 mg/ml to 0.2 mg/ml and first investigated its effect on mitochondrial respiration in *w^1118^*, *w^1118^*-mt237, ORT and ORT-mt237 3^rd^ instar larvae. In general, 0.2 mg/ml exposure of CA decreased the respiration of each tested fly line, but this effect was significant only in ORT line (**Fig. S2**). Next, we examined whether 0.2 mg/ml CA is sufficient to induce the formation of melanotic nodules in ORT cybrid lines mt237 and mt765 and simultaneously checked the sex of the dissected larvae. Also this lower concentration was effective in inducing melanotic nodules, and females were shown to be overrepresented among the nodule positive larvae (**Fig. 4A**). To examine the transmissibility of the CA induced nodule phenotype, we reared larvae on control diet or food supplemented with 0.2 mg/ml CA. Eclosed adults were collected, sexed and kept 3-5 days in control food before performing the following crossing scheme using virgin flies: a) control females x control males, b) control females x CA treated males, c) CA treated females x control males, d) CA females x CA treated males. The highest nodule incidence occurred in offspring of parents where the unexposed control females were crossed with the CA exposed males. The same result was observed in both study lines. Importantly, the study design shows that the nodule phenotype can be transmitted to the following generation, mainly via the CA exposed paternal lineage (**Fig. 4B**).

**Figure 4.**
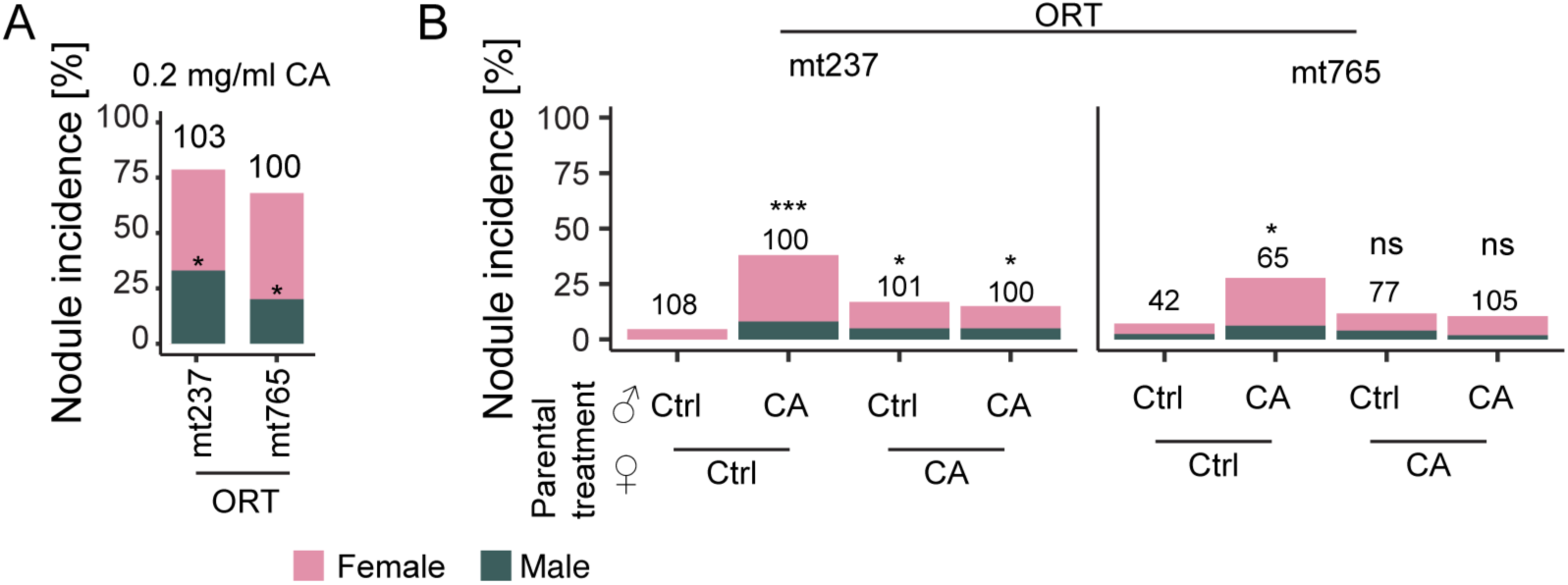
CA-induced aberrant immune activation is female biased and transmitted to the next generation via paternal lineage. **A)** Percentage of nodule-positive larvae following rearing on food supplemented with 0.2 mg/ ml CA, showing the proportions of females and males within the phenotype. Asterisks indicate statistically significant differences between female and male nodule frequencies. The number of dissected larvae is indicated in the plot. **B)** Transgenerational inheritance of the 0.2 mg/ml CA-induced melanotic nodule phenotype in the offspring of parents treated at the larval stage. Melanotic nodule phenotype was inspected from the offspring of the following crosses: control female x control male, control female x CA male, CA female x control male and CA female x CA male. The number of dissected individuals is indicated in the plot. Asterisks indicate the statistical difference of each combination of parental treatment to the control (progeny of the untreated control cross). ns, not significant; * p< 0.05; ** p< 0.01; *** p< 0.001.

**Figure 5.**
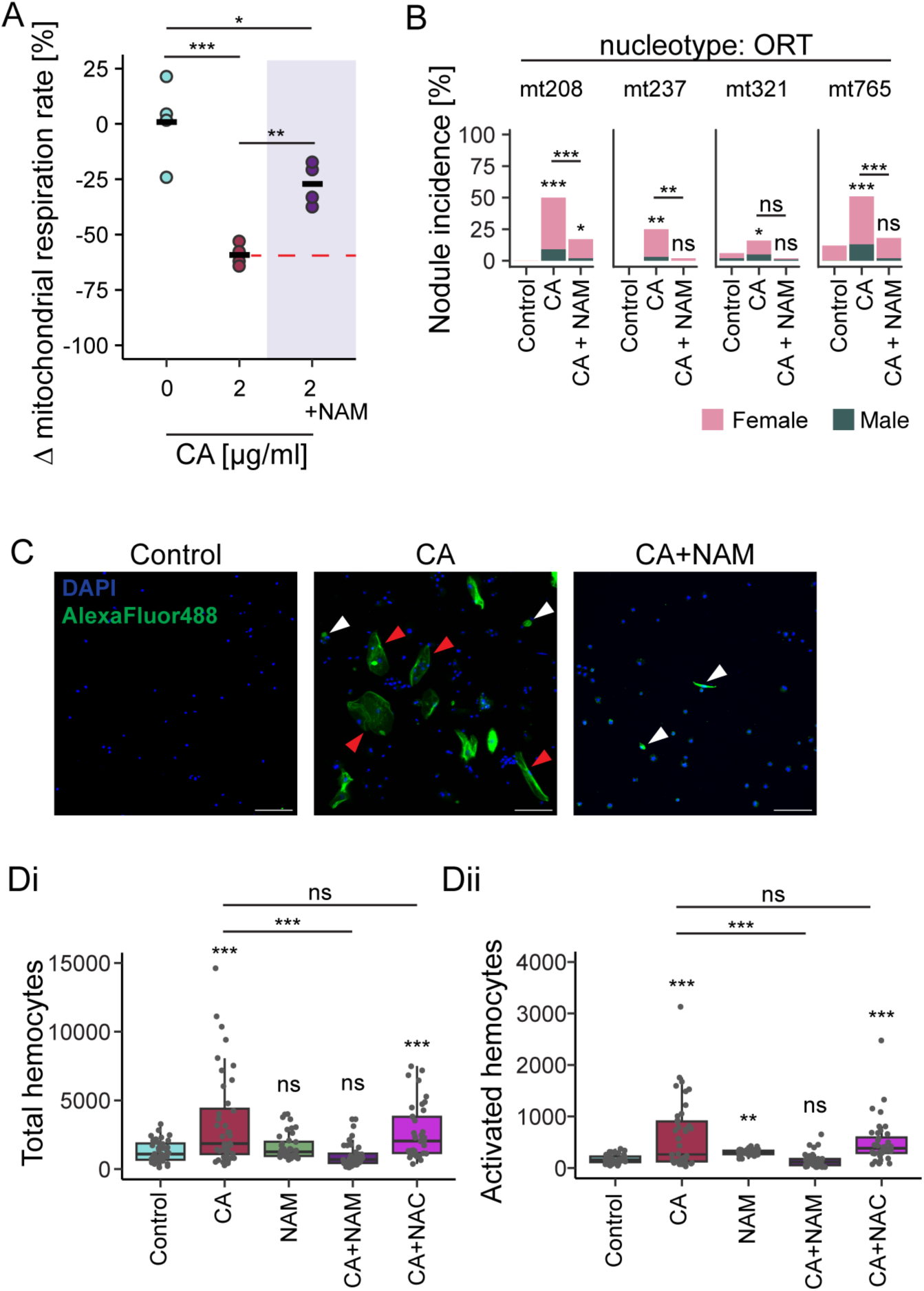
Co-supplementation of nicotinamide (NAM) rescues CA-induced decrease in mitochondrial respiration and the induction of aberrant immune activation. A) Oxygen consumption of ORT larvae reared on control food, or that supplemented with CA (2 mg/ml) or CA + NAM (300 µg/ml), normalized to the respiration of control larvae. B) Percentages of nodule positive 3^rd^ instar larvae after rearing them on control, CA or CA + NAM supplemented food. A total of 100 3^rd^ instar larvae were dissected for each group, except for the mt321 and mt765 controls groups, for which 121 and 125 larvae were dissected, respectively. Asterisks on top of each bar refer to statistical significance compared to the control. C) Immunostaining of hemocytes collected from ORT-mt765 3^rd^ instar larvae reared with control diet, or diet supplemented with 2 mg /ml CA or 2 mg/ml CA + 300 µg/ml NAM. Images show fixed hemocytes with DAPI nuclear stain (blue) and L2 lamellocyte antibody (AlexaFluor488nm, green). Examples of L2-positive lamellocytes and lamellocyte aggregates are marked with red arrowheads, L2-positive hemocytes with non-lamellocyte morphology with white arrowheads. D) Nodule-prone ORT-cybrid (nORT-mt237) larvae were reared on control diet or those supplemented with 2 mg/ml CA, 300 μg/ml NAM, CA + NAM, or CA + 1 mg/ml N-acetyl cysteine (NAC). Hemocytes from 3^rd^ instar larvae were subsequently profiled using flow cytometry to measure **Di)** total hemocyte counts and **Dii)** activated hemocyte counts. Asterisks on top of each boxplot refer to statistical significance compared to the control. ns, not significant; * p< 0.05; ** p< 0.01; *** p< 0.001.

### Nicotinamide supplementation rescues the effects of CA on decreased mitochondrial respiration and aberrant immune activation *in vivo*

Based on the metabolome data, B_3_ vitamers NAM and niacin were depleted in adult flies after rearing the larval stages in CA supplemented food (**Fig. 2Cii**). We investigated whether co-supplementation with CA and NAM could mitigate the CA-induced decrease in mitochondrial respiration. We reared ORT larvae on control diet and on food supplemented with 2 mg/ml CA or CA + 300 µg/ml NAM. Co-supplementing NAM significantly enhanced the mitochondrial respiration of the larvae (**Fig. 5A**). As NAM supplementation alleviated the CA-induced reduction in mitochondrial respiration, we next examined whether NAM could also attenuate the aberrant immune activation induced by CA. Co-supplementation with NAM led to a rescue effect in the examined nodule prone ORT cybrid lines, as it reduced the incidence of melanotic nodules to the control level (**Fig. 5B**), and the presence of immune activated hemocytes shown with fluorescent imaging (**Fig. 5C, Fig. S1A**).

To determine whether the ability of NAM to counteract the CA induced aberrant immune activation is related to its antioxidant effects^86,87^, we supplemented the *Drosophila* food with CA only, NAM only, CA + NAM and CA with the generic antioxidant N-acetylcysteine (NAC, 1mg/ml), which acts via the glutathione system. We have earlier shown that NAC reduces ROS when fed to the larvae^41^. Profiling the hemocytes with flow cytometry showed that NAM counteracts the CA-mediated expansion of hemocytes in general, and those of lamellocytes, whereas CA + NAC hemocyte profiles resembled those of CA treated larvae (**Fig. 5D**). Based on these findings the rescue effect of NAM is not dependent on its antioxidant effect.

To assess the therapeutic potential of combined CA and NAM/niacin supplementation during bacterial infection, and to rule out any antagonistic effects on CA’s antibacterial activity, we examined the effects of CA and NAM/niacin, individually or in combination, on the growth of Gram-positive bacterium *Staphylococcus aureus* and Gram-negative bacterium *Escherichia coli*, by evaluating their bacteriostatic/bactericidal effects. We show that CA alone, CA+NAM and CA+niacin, in concentrations matching the experiments carried out with HEK293T cells, inhibit the growth of *S. aureus* and *E. coli*, indicating that NAM or niacin do not interfere with the bacteriostatic action of CA (**Fig. S3**). Furthermore, NAM or niacin alone do not have bactericidal or bacteriostatic activities (**Fig. S3**).

### Larvae with melanotic nodules have altered levels of kynurenine pathway metabolites

We have previously demonstrated that hemocyte-targeted OXPHOS gene knockdowns in *D. melanogaster* larvae trigger hemocyte activation and the formation of melanotic nodules^41^. Here, we investigated whether NAM treatment could alter the nodule phenotype of these larvae. For this, we selected the OXPHOS complex V gene *ATPsynCF6* (*ATP synthase, coupling factor 6*) knockdown line that manifested one of the highest melanotic nodule incidence among the previously tested OXPHOS knockdown lines^41^, and shows a robust gene silencing in hemocytes (**Fig. 6A**). However, hemocyte-targeted *ATPsynCF6* knockdown larvae reared on NAM-supplemented food exhibited a nodule phenotype similar to that of larvae reared on the control diet (**Fig. 6B**), indicating that NAM supplementation does not rescue the nodule phenotype when OXPHOS perturbation is induced genetically and hemocyte-specifically. Another feature distinguishing this model from the CA-induced nodule phenotype was the lack of female predominance among nodule-positive larvae (**Fig. 6B**).

**Figure 6.**
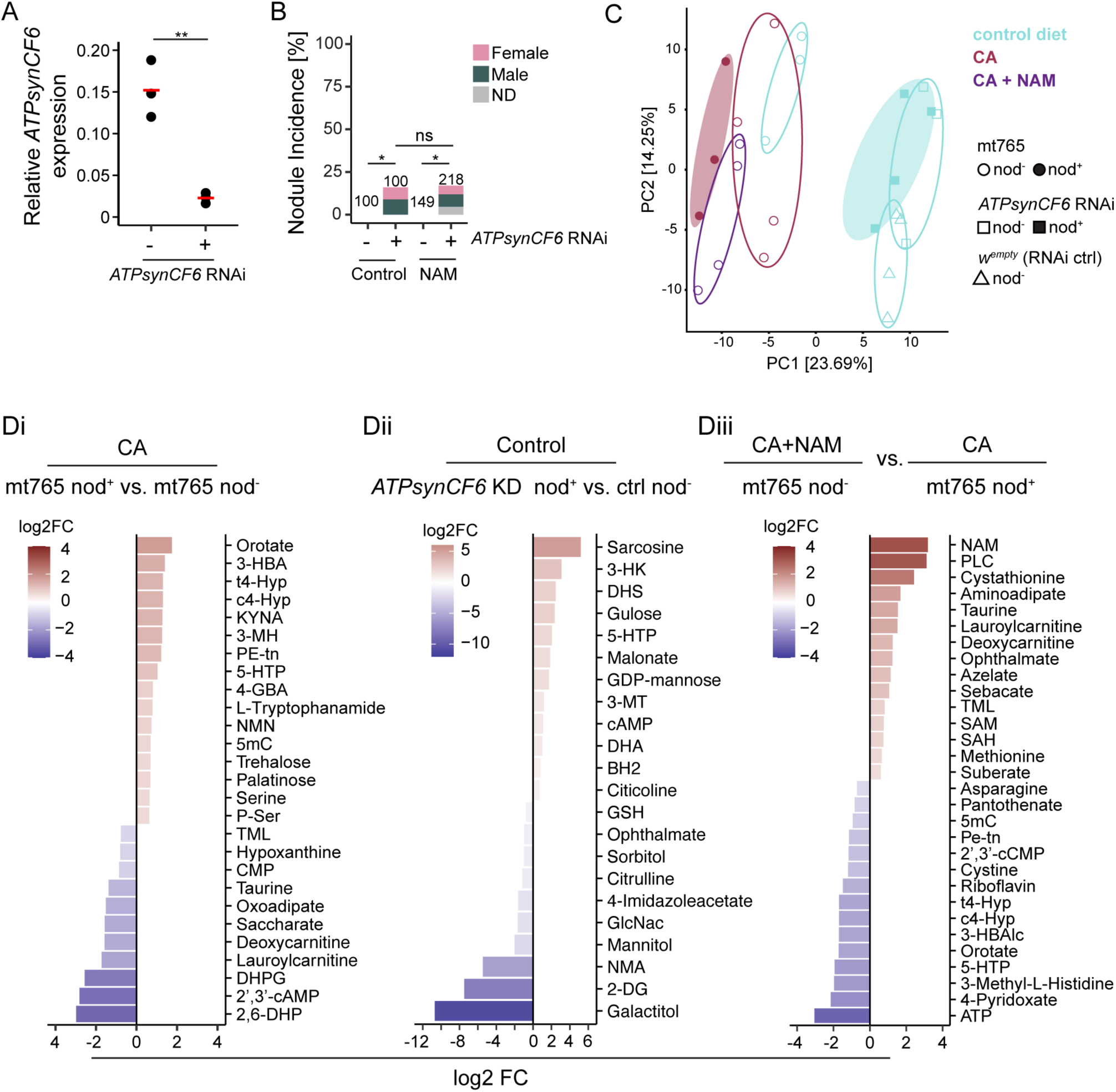
CA-induced aberrant immune activation is connected to altered tryptophan metabolism. **A)** Knockdown efficiency of the hemocyte targeted *ATPsynCF6* RNAi line was measured with qRT-PCR from the hemocytes collected from 3^rd^ instar larvae in comparison to the control (w^empty^) with normal *ATPsynCF6* expression. Data were analyzed using a one-way Analysis of Variance. **B)** Nodule percentages of larvae with normal level of *ATPsynCF6* gene expression and hemocyte targeted *ATPsynCF6* knockdown larvae reared on control diet or that supplemented with 300 µg/ml NAM. The number of dissected larvae is indicated in the plot. **C)** Principal component analysis (PCA) performed with the 3^rd^ instar larval samples reared on control diet, that supplemented with 2 mg/ml CA, or 2 mg/ml CA + 300 µg/ml NAM. Nod^+^ and nod^-^ larvae were included from the nORT mt765 and *HH> ATPsynCF6* larvae and nod^-^ larvae from ORT and *HH>w^empty^*. **D)** Metabolome analysis of the larval samples showing significantly increased or reduced metabolites (max. 30) when comparing **Di)** CA exposed ORT-mt765 nod^+^ larvae to their nod^-^ counterparts **Dii)** control food reared immune cell targeted *ATPsynCF6* RNAi nod^+^ larvae with their nod^-^ counterparts, and **Diii)** CA + NAM exposed ORT-mt765 nod^-^ larvae with CA exposed nod*^+^* counterparts. ns, not significant; * p< 0.05; ** p< 0.01; *** p< 0.001.

We performed another set of metabolome analyses with 3^rd^ instar larvae to study further which metabolites and pathways are affected by CA, CA + NAM and those connected to the nodule phenotype. For this, we selected the ORT and nodule prone ORT-mt765 cybrid line, and the genetically induced nodule model, *ATPsynCF6* RNAi in hemocytes, and its respective control (genetic background of RNAi line). We sampled ORT and mt765 reared with control diet, CA supplemented diet and that co-supplemented with CA + NAM. From the CA treatment we included mt765 larvae that had nodules (+) and those without (-) nodules and additionally included +/- nodule bearing larvae from the *ATPsynCF6* RNAi line reared in control food. Based on the PCA analysis of the nodule-producing genotypes (mt765 and *ATPsynCF6* knockdown; **Fig. 6C**), stronger clustering occurred based on the genetic background (PCA1; 23.7%) than on the treatment (PCA2; 14.3%). Among the mt765 samples, CA-treated nodule-bearing (nod^+^) larvae formed a distinct cluster separate from CA-treated larvae that lacked nodules (nod^-^), whereas the CA + NAM samples clustered closed to the control samples (**Fig. 6C**). Following the *ATPsynCF6* knockdown, nod+ and nod^-^ samples were overlapping, while the corresponding control sample was more separated (**Fig. 6C**).

When examining the metabolites that were increased or decreased following CA treatment in ORT and mt765 (nod^−^) relative to their respective untreated controls, the two lines exhibited mostly similar responses. Both lines had elevated levels of metabolites related to oxidative stress (GSH, taurine and hypotaurine in ORT; GSH and ophthalmate in mt765) and depletion of microbiota-associated metabolites (homoserine, 4-Imidazoleacetate, hippurate, imidazoleacetate, hydroxyphenyllactate, hydroxyphenyllactic acid, 3-(2-hydroxyphenyl)propanoate, homovanillate, N-methylglutamate and 4-hydroxybenzoate in ORT and 4-Imidazoleacetate, and 3,4-dihydroxybenzoate in mt765) (**SFig. 4Ai** and **Aii**). Furthermore, CA-exposed mt765 nod^-^ larvae showed elevated levels of metabolites related to tyrosine/catecholamine metabolism (homogentisate, 3,4-dihydroxyphenylglycol, and 4-methylcatechol) and tryptophan metabolism (N-acetylserotonin and xanthurenate). In addition, CA treatment was associated with remodeling of NAD metabolism, characterized by reduced nicotinamide (NAM) and NADH and increased nicotinamide mononucleotide (NMN) (**Fig. S4Aii**).

The PCA plot (**Fig. 6C**) indicated that the formation of melanotic nodules is connected to specific metabolites that distinguish these larvae from other CA treated larvae, and that co-supplementation with NAM rescues the effects connected to nodules at the metabolite levels. Hence, we next checked the significantly up-and down regulated metabolites among the CA treated nod^+^ vs nod^-^ ORT-mt765 larvae (**Fig. 6Di**). Metabolites that were more abundant in nod^+^ larvae compared to nod^-^, indicated the involvement of the tryptophan pathway, involving the metabolites 5-hydroxy-L-tryptophan (5-HTP), kynurenic acid (KYNA) and indirectly serine, which is a cofactor in tryptophan metabolism (**Table S3**). The tryptophan pathway is a metabolic branch point where the cell produces kynurenine metabolites (such as kynurenine, 3-HK and kynurenic acid), or melatonin via serotonin pathway. In the nod^+^ larvae, metabolism seems to shift towards the kynurenine pathway. In mammals, kynurenine pathway is the predominant route associated with immune regulation and NAD^+^ biosynthesis^88,89^, whereas serotonin-melatonin pathway contributes to circadian regulation and antioxidant defense^90,91^. Moreover, active kynurenine pathway has been shown to impair mitochondrial β-oxidation, and kynurenine-associated oxidative stress impairs fatty acid oxidation and promotes metabolic dysfunction, whereas melatonin supports efficient β-oxidation^92^. In *Drosophila*, kynurenine pathway likely does not contribute to *de novo*NAD^+^ synthesis due to lack of some of the key enzymes leading to NAD^+^. Among the decreased metabolites in nod^+^ larvae were deoxycarnitine and lauroylcarnitine (involved in mitochondrial fatty acid transport), and oxoadipate, all associated with mitochondrial β-oxidation. Furthermore, when nodule-bearing *ATPsynCF6* RNAi larvae were compared to their controls, metabolites of the kynurenine pathway were among the most abundant ones (**Fig. 6Dii**, **Table S3**), including 3-HK and 5-HTP.

Importantly, several metabolites altered in CA-exposed mt765 nod⁺ larvae were restored by CA + NAM co-supplementation in mt765 nod⁻ larvae, including NAM itself. Taurine, TML, deoxycarnitine, and lauroylcarnitine were decreased in CA-treated mt765 nod⁺ larvae and upregulated after NAM treatment (**Fig. 6Diii**). Other metabolites that were up in mt765 nod^+^ larvae and rescued with NAM included orotate, trans-4-Hydroxyproline (t4-Hyp), cis-4-Hydroxyproline (c4-Hyp), Phosphoethanolamine (PE-tn), 5-HTP and 5-Methylcytosine (5mC), being consistent with normalization of stress-associated metabolic programming.

### The CA induced defect in respiration and mitochondrial thermogenesis is rescued by vitamin B_3_ co-supplementation in a human cell line

Co-supplementation with NAM rescued the CA-caused reduction in mitochondrial respiration (**Fig. 5A**) and the immune activation in *Drosophila* cybrid larvae (**Fig. 5B-D**). Next, we sought to validate the *in vivo Drosophila* findings regarding the adverse effects of CA on mitochondrial respiration and thermogenesis using a human cell line model, and to determine whether the vitamin B_3_ co-supplementation strategy could alleviate the adverse effects of CA also in a mammalian model. HEK293T cells, a human embryonic kidney cell line widely used in mitochondrial research^93^, were exposed to 100 μg/ml CA and cell viability was assessed at 17, 24, 30, 48, 72 and 96 hours following treatment, as well as in untreated control cells. After 17 and 24 h exposure to CA, cell viability remained at approximately 90% in the control group, whereas it was around 75% in the CA-treated group. At later time points, viability declined markedly in both groups (**Fig. 7A**). Based on the viability assay, a 24-hour exposure period was selected, and the HEK293T cells were exposed to 20 or 100 μg/ml CA. The effects on mitochondrial respiration were assessed, showing that the lower dose caused a modest but significant reduction in mitochondrial respiration, whereas the higher dose reduced oxygen consumption by an average of 70% (**Fig. 7Bi**).

**Figure 7.**
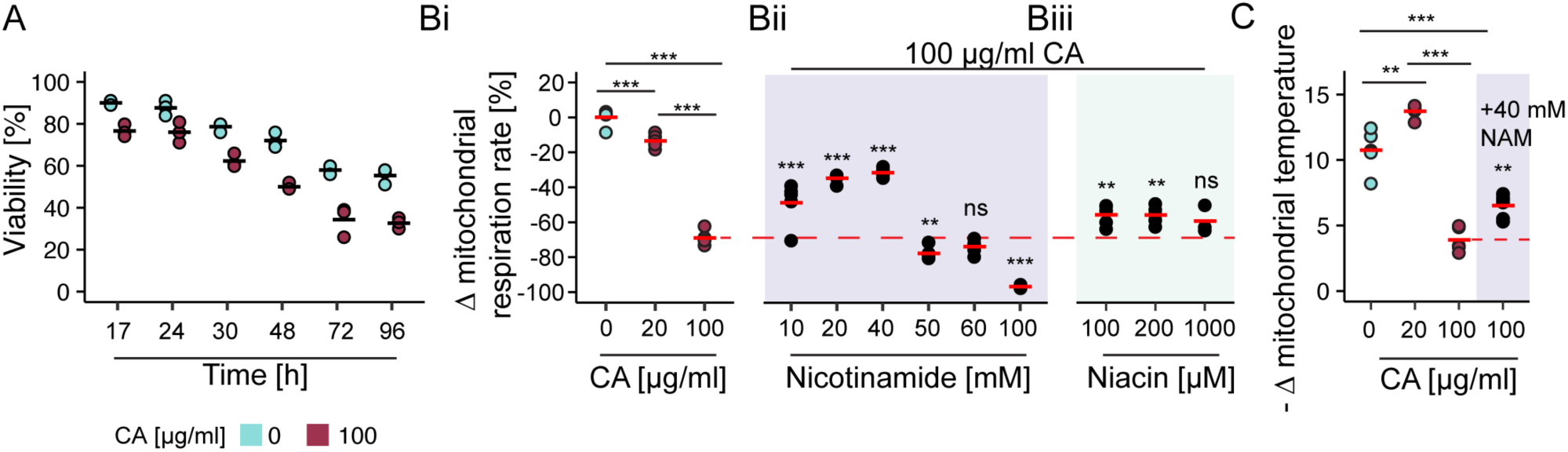
Vitamin B_3_ vitamers alleviate the CA-induced decrease in mitochondrial respiration and thermogenesis in a human cell line. **A)** Viability of HEK293T cells following exposure to 100 µg/ml CA, expressed relative to time-matched untreated controls and normalized to the viability of control cells at 0 h. Three replicates per treatment were used. **B)** Mitochondrial respiration (oxygen consumption) rates of HEK293T cells after supplementation with **Bi)** low and high doses of CA, and co-supplementing 100 µg/ml CA with several doses of **Bii)** NAM (light purple background) or **Biii)** niacin (light green background). Red dashed line shows the rescue effect of co-supplementing CA with NAM or niacin when compared to the 100 µg/ml CA treated cells. Five replicates were used for each treatment except for 100 mM nicotinamide (4 replicates) and 1000 μM niacin (3 replicates). **C)** Mitochondrial temperature was measured with MTY from the unexposed HEK293T cells and those exposed for 24 hours to 20 μg/ml or 100 μg/ml CA and after co-supplementing CA (100 μg/ml) with NAM (40 mM; purple background). Data were obtained from five to six replicates. ns, not significant; * p< 0.05; ** p< 0.01; *** p< 0.001.

Next, we evaluated the effects of increasing concentrations of the vitamin B_3_ vitamers NAM and niacin on the oxygen consumption rates of otherwise untreated HEK293T cells. 24-hour exposure to 10, 20, 40 and 100 mM NAM showed that while the lower concentrations had no effect on the respiration, 40 and 100 mM significantly decreased the respiration (**Fig. S5**). The effect of 100, 200 and 1000 uM of niacin exposure showed that the lower doses had no effect on the respiration, while the highest dose significantly increased it (**Fig. S5**).

We then co-supplemented the HEK293T cells with 100 μg/ml CA, which was shown to decrease the mitochondrial respiration, with selected concentrations of NAM or niacin to determine whether they could mitigate the CA-induced impairment of mitochondrial respiration. Co-supplementation with the lowest dose of NAM, 10 mM, already significantly enhanced the mitochondrial respiration of 100 μg/ml CA exposed cells (**Fig. 7Bii**). Respiration was gradually enhanced with the higher doses 20 and 40 mM of NAM, until not alleviating the CA effects or decreasing the respiration with the doses 50, 60 and 100 mM (**Fig. 7Bii**). Co-supplementation of niacin, however, did not manifest a strong dose-response curve, as 100 and 200 μM concentrations provided modest alleviation on the respiration of CA exposed cells, whereas oxygen consumption rates after exposure to 1000 μM niacin started to resemble that of the CA treated cells (**Fig. 7Biii**).

We then assessed the effects of the low (20 μg/ml) and high (100 μg/ml) CA concentrations on mitochondrial thermogenesis in HEK293T cells. The lower dose caused a mitohormetic effect, with a modest but consistent increase in the mitochondrial temperature (**Fig. 7B**). With the higher dose, we observed a marked decrease in mitochondrial temperature (**Fig. 7B**). Notably, co-supplementation with 40mM NAM, which produced the strongest rescue of the respiratory phenotype, significantly restored mitochondrial heat generation in cells exposed to 100 μg/ml CA (**Fig. 7B**).

Taken together, these findings highlight the utility of the *Drosophila* model for investigating mitochondrial dysfunction and demonstrate the evolutionary conservation of the effects of CA on mitochondrial function, as well as the ability of vitamin B_3_ vitamers to alleviate these defects. The concordant findings observed in both human cell line and the *Drosophila in vivo* model further support the relevance and translational potential of the identified *Drosophila* phenotypes and underlying mechanisms to mammalian systems.

## Discussion

We modelled the adverse effects of chloramphenicol (CA) in *D. melanogaster*, focusing on dose-dependent responses, potential immunomodulatory effects and the role of host genetic background with the aim of identifying strategies to alleviate CA-induced mitochondrial stress in human cell line model. We confirmed that CA represses mitochondrial respiration in *Drosophila* derived S2 cell line and in *Drosophila* larvae, in accordance with other *in vivo* and *in vitro* models ^20,94^, in a dose-dependent manner. *In vivo*, the selected concentrations of CA, fed to the *Drosophila* larvae throughout their development, caused only modest developmental delay. We also demonstrate that, in addition to effects on respiration, also mitochondrial temperature responds to CA exposure in dose-dependent manner in human and *Drosophila* cell lines. To our knowledge, no previous studies have examined the impact of mitochondria-affecting antibiotics on mitochondrial thermogenesis. One of the most well-documented CA caused disease is the gray baby syndrome, caused by CA accumulation due to immature hepatic metabolism in neonates. Gray baby syndrome is characterized by several symptoms, including cyanosis, circulatory collapse and hypothermia^95^. Based on our findings, the hypothermia could result from decrease in mitochondrial temperature at the systemic level caused by exposure to CA.

In both insect and mammalian models, CA induced a mitohormetic response, with low-dose exposure causing a mild impairment of mitochondrial respiration accompanied by increased mitochondrial heat generation, whereas higher CA concentrations led to a substantial decline in both respiration and thermogenesis. In mitohormesis, a mild mitochondrial stress initiates coordinated metabolic and cellular adaptations that improve the cell’s capacity to withstand subsequent environmental or physiological stressors^96^. We have earlier shown that mild mitochondrial perturbations, caused by natural mtDNA variation or experimentally induced genetic mitochondrial insults can result in aberrant immune activation and enhance the immunocompetence of the host when encountering bacterial, viral or parasitotic pathogens^29,41^. This enhancement of immunocompetence was connected to mitochondrial unfolded protein response (UPR^mt^)^41^, which is a cellular stress response pathway and activated when misfolded or damaged proteins accumulate inside mitochondria, to maintain mitochondrial function under stress conditions^97^. As CA affects the translation of the mtDNA encoded proteins and causes mitochondrial stress, we were interested to observe its possible effects on immunity. Furthermore, as there is evidence that specific antibiotics can cause adverse and permanent effects when given to a host carrying certain mtDNA mutations, such as in the case of aminoglycoside-induced deafness caused by the destruction of sensory hair cells^98^, we studied the effects of CA exposure with naturally occurring mtDNA variants known to possess variation on their rRNA genes. These *Drosophila* cybrid lines that carry mitochondrial rRNA (*16S* or *12S*) variants develop normally when maintained on control food, suggesting that the defining variants are largely benign under control conditions. However, the cybrids of both nuclear backgrounds were more affected by the CA exposure as they generally experienced a developmental delay on CA supplemented food. We have not examined the structural interactions between these variants and CA, but it is unlikely that all the variants (**Table 1**) affect CA interaction in the same manner. Additionally, we did not observe major differences in basal respiration or in respiration following CA exposure between the background lines and the studied cybrid lines.

Upon CA exposure, activation of cell-mediated innate immunity was observed only in ORT cybrids, as indicated by immune cell activation (lamellocyte differentiation) and the formation of melanotic nodules. This background specific response is consistent with previous evidence showing that *w^1118^* and ORT differ in their susceptibility to mitoribosomal stress. Jacobs *et al.*^99^ showed that the *tko^25t^* mutation affecting mitochondrial ribosomal protein S12 (mRpS12) caused severe mitochondrial dysfunction associated phenotypes in the ORT background, whereas the same mutation produced only moderate developmental delay and relatively mild bang sensitivity phenotype (transient seizure/paralysis upon agitation) in *w^1118^*. Their finding also indicates that the ORT background appears more prone to mitoribosomal stress and that genetic modifiers affect mitochondrial stress responses between the two studied nuclear backgrounds.

The melanization reaction leading to melanotic nodules can be triggered in both humoral and cellular responses and best characterized as part of the *Drosophila* host defense against wasp parasitation, but has been described as an outcome of >100 genetic mutations^36^ and various experimental conditions^29,41,55,82^. The CA-induced immune phenotype strengthens the proposed link between mitochondrial stress and immune activation. Our previous studies have demonstrated that melanotic nodules can be induced by mitochondrial stress either through (i) dietary exposure to antimycin A^82^ an inhibitor of OXPHOS complex III^82^, (ii) naturally occurring mtDNA mutation in OXPHOS complex III gene *CYTB*^29^ and iii) hemocyte-specific knockdown of OXPHOS subunits^41^. In a separate study, ciprofloxacin, an antibiotic with mitochondrial toxicity but a non-translational mechanism of action, was reported to induce the formation of “black spots” in *w^1118^* larvae^100^. Although this phenotype was not investigated further, it is likely that these black spots represent melanotic nodules. However, CA-induced nodules clearly differ from those induced by hemocyte-specific knockdown of OXPHOS complex v gene *ATPsynCF6,* as the nodule phenotype in CA exposed larvae is strongly biased towards females and are counteracted by co-supplementing vitamin B_3_ vitamer nicotinamide. Despite of these differences, they both shared metabolic features involving altered tryptophan catabolism through the kynurenine pathway. However, changes in the tryptophan pathway can be considered primarily as a marker of metabolic and mitochondrial stress associated with immune activation, rather than as direct drivers of melanotic nodules.

The widely used *Drosophila* line *w^1118^* carries a loss-of-function mutation in the *white* (*w*) gene encoding for a transporter involved in transport or pigment precursors (tryptophan, guanine and kynurenine) and resulting in, among other issues, the characteristic white-eye phenotype. This mutation leads to altered kynurenine metabolism, associated with changes in oxidative stress tolerance and stress responses^101^. Because several kynurenine-pathway intermediates, like 3-HK, are redox-active, alterations in this pathway could potentially influence physiological processes sensitive to redox balance, including melanization. This could partially explain some of the differences observed between the CA caused effects in *w*^111^*^8^* and ORT lines.

Differences between females and males in immunity are consistently observed in mammals, and females typically mount stronger innate and adaptive immune responses than males^102^. However, this female advantage is accompanied by an increased susceptibility to inflammatory and autoimmune diseases^102^. Sex differences in *Drosophila* immunity have been investigated predominantly in the context of humoral innate immune responses to bacterial and viral infections^103^. Although melanotic nodule phenotypes have been widely reported in *Drosophila*, to our knowledge no sex-bias has previously been reported and there are only limited work examining sex-specific responses to parasitoid infections^104^ or cell-mediated immunity^105^. Although our data indicates that the CA induced nodule phenotype can be transmitted to the next generation via paternal lineage, the nodule phenotype itself is female biased. These results show that antibiotics, especially those affecting mitochondria, might have far-reaching sex-specific consequences that should be avoided by alleviating the mitochondrial stress induced by antibiotics.

Two general effects of CA on metabolite profiles emerged from metabolomic analyses across genotypes and developmental stages: impaired mitochondrial β-oxidation and carnitine biosynthesis, and vitamin B_3_ metabolism. While CA is well known to inhibit mitochondrial protein synthesis, its effects on fatty-acid β-oxidation have received considerably less attention. Given the dependence of β-oxidation of NAD^+^ and FAD generated by ETC, impaired due to CA treatment, our metabolomic findings suggests that CA-induced mitochondrial dysfunction may extend to lipid catabolism, resulting in reduced mitochondrial fatty acid oxidation, and promoting diversion of fatty acids toward alternative catabolic pathways such as ω-oxidation^106^. The reduction of carnitine intermediates is more surprising, but may reflect reduced mitochondrial fatty-acid flux secondary to impaired ETC.

Vitamin B_3_ vitamers nicotinamide (niacinamide), nicotinamide riboside, nicotinamide mononucleotide and niacin (nicotinic acid) act as precursors for NAD^+^, raising the possibility that CA treatment affects NAD^+^ metabolism. It is known that deficiency of niacin or tryptophan causes pellagra (dermatitis, diarrhea, dementia, death), and CA has been reported among drugs associated with secondary pellagra in humans^107^. Long-term CA use may also induce hematotoxicity, manifested as bone marrow suppression and aplastic anemia, via mechanisms of which are still unclear. It has been speculated that CA treatment could activate T-lymphocytes, which could kill hematopoietic cells or that impaired mitochondrial function disrupts erythrocyte maturation^18^. However, another plausible explanation could be their heightened sensitivity to vitamin B deficiencies^108^.

The effects of vitamin B_3_ vitamers NAM and niacin were studied further in our study. NAM is converted to NAD^+^ through the salvage pathway, whereas niacin is converted through the Preiss-Handler pathway^109^. The two vitamers additionally differ in their chemistry and biological effects, including niacin’s effect on blood lipids^74^. In this study, the outcomes associated with NAM supplementation were more promising than those observed with niacin, as NAM rescued the CA caused mitochondrial perturbations more efficiently than niacin. Co-supplementing NAM rescued the CA effects on respiration both in *Drosophila* larvae and in HEK293T cells and rescued the adverse effects of CA on the mitochondrial heat generation in HEK293T cells. Notably higher doses of NAM (50 mM NAM with CA and >40 mM NAM in untreated cells) reduced the mitochondrial respiratory capacity in HEK293T cells. Although generally well tolerated, detrimental effects of higher doses of NAM have been reported, including inhibition of sirtuin activity and consequent alterations of mitochondrial metabolism^110,111^.

These observations highlight the importance of correct dosing of mitochondrial boosters. To further validate the CA effects and the effect of co-supplementing NAM or niacin, other cells lines could be recruited, such as human derived neuronal-(SH-SY5Y), liver-(HepG2) or skeletal muscle myoblast (LHCN-M2) cell lines.

Notably, all CA effects observed in this study (respiration, thermogenesis, aberrant immune activation) were consistently mitigated by vitamin B_3_ co-supplementation across all experimental settings. As CA directly inhibits mitochondrial protein synthesis, it can disrupt mitochondrial function and increase the cellular demand for NAD(H)-related metabolism, potentially contributing to a functional vitamin B_3_ deficiency. In addition, as a broad-spectrum antibiotic, CA may alter the host microbiota, and several intestinal bacterial species possess pathways to synthesize vitamin B_3_-related compounds, including NAM and niacin, contributing to the host NAD^+^ pool^112^. Consequently, CA exposure could affect host vitamin B homeostasis through both direct mitochondrial effects and indirect microbiota-mediated mechanisms. As the commensal microbiota was not assessed before or after CA treatment, potential differences between the *w*^111^*^8^* and ORT background lines and their corresponding cybrid lines cannot be excluded. Hence, whether these putative microbiota differences influenced the aberrant cell mediated immune activation and the development of the nodule phenotype, remains unknown.

The effect of CA and vitamin B_3_ supplementation on mitochondrial respiration and thermogenesis in HEK293T cells were measured 24 hours after the onset of treatment, and the impact of longer exposure times was not investigated. However, in our *in vivo* model, CA and NAM were administrated to *Drosophila* through their larval development within their diet, providing a longer-term, physiologically relevant route of exposure, and enabling systemic exposure of the whole organism. Furthermore, samples collected at two different developmental stages, both during and following CA treatment, exhibited similar metabolomic profiles, suggesting that the effects of CA exposure persist beyond the treatment period.

These results provide a rationale for exploring vitamin B_3_ (specifically NAM) supplementation as an adjunctive approach to reduce CA-associated toxicity and potentially expand its clinical utility in severe infections for which alternative antibiotic options are limited. Furthermore, our work demonstrates that *Drosophila* is not only a powerful genetic model organism but also an informative platform for pharmacological and toxicological research, enabling the efficient identification of drug-induced adverse effects and potential therapeutic interventions.

## Acknowledgements

We thank Yuliya Basikhina, Juha Koivuniemi, Mahmudul Tanvir, Pihla Tiilikainen and Milla Pihlajamäki for their assistance in various experiments. We further thank Nick Lane, Brendan Battersby and Jaakko Pohjoismäki for their valuable comments regarding the doctoral thesis of Tilman Tietz. The *Drosophila* work was carried out at the Tampere *Drosophila* Facility (Tampere University) which is partly funded by Biocenter Finland. We further acknowledge Imaging Facility, Protein Services, and Cellular Respiration and Energetic Metabolism facility of Tampere University.

## Data availability

All the data presented in this paper are included in the Figures and Supplementary Tables.

## Author contributions

Designed and conceived the project: TSS, LV, TTi, MT

Performed data generation and analyses: TTi, MT, TTu, LV, KMa, KK, TP, PL, EN, KL, SV, TSS

Contributed data or analysis tools: TTi, LV, JJ, K-ML, KMo, VH

Contributed to discussion and interpretation: TSS, LV, TTi

Wrote the paper; all authors contributed to the editing process. TSS, TTi, LV

## Supplementary figures and figure captions

### List of Supplementary Tables

**Table S1.** Data for respirometry experiments in figures 1A-B, 5A, 7Bi-Biii, S2 and S5, and for mitochondrial thermometry Figures 1Aii and 7C.

**Table S2.** Data for figures 2Ai-Aii, 3A, 3C, 3E-F, 4A-B, 5B, 5Di-Dii, 6A-B, S1.

**Table S3.** Metabolite fold changes and p-values for figures 2Ci-Cii, 6Di-Diii, S4B-C **Table S4.** Raw and normalized peak values for metabolite data.

---

**Figure S1.**
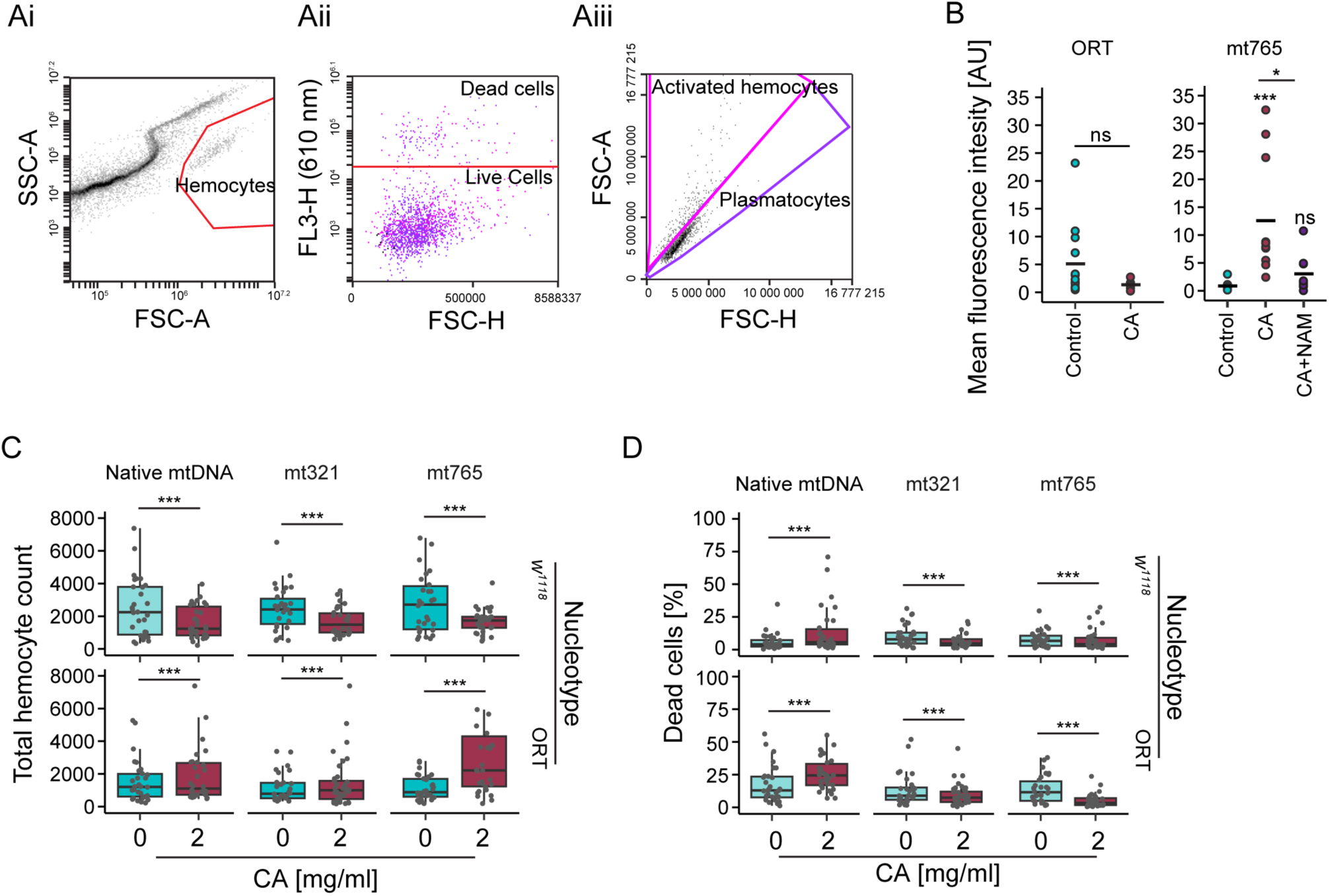
L2 antibody staining quantification, hemocyte gating strategy, total hemocyte counts and hemocyte viability following CA-exposure. Ai-Aiii) Flow cytometry gating strategy for *Drosophila* larval hemocytes. Ai) Hemocytes were separated from debris based on their size (FSC-A) and granularity (SSC-A). Aii) PI-negative (live) and positive (dead) hemocytes were separated based on FL3-H axis (610 nm), and Aiii) live hemocytes were further separated based on their shape on FSC-A vs FSC-H plot to distinguish between plasmatocytes (round cells) and activated hemocytes (mainly lamellocytes, irregular shape). B) Quantification of the AlexaFluor 488nm fluorescence intensity from hemocytes of control and 2 mg/ml CA fed ORT and mt765 larvae and mt765 larvae fed with CA (2 mg/ml) + NAM (300 µg/ml). Each dot represents a microscopy image (examples given in Fig. 3D and 5C). Total cell counts in each treatment was: ORT control 513, ORT CA 192, mt765 control 194, mt765 CA 1394, mt765 CA + NAM 196. Asterisks on top of treatments refer to statistical significance compared to the control. C) Total number of hemocytes of 3^rd^ instar larvae reared either on control food or food supplemented with 2mg /ml CA. D) Percentage of dead cells based on the PI-staining. ns, not significant; * p< 0.05; ** p< 0.01; *** p< 0.001.

**Figure S2.**
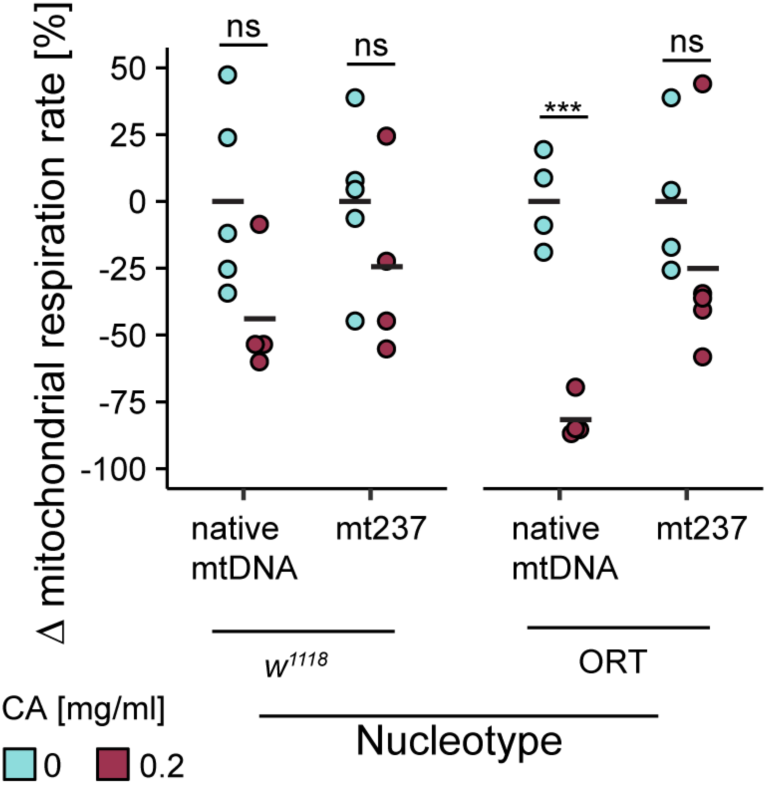
CA exposure causes genotype specific effects on mitochondrial respiration. Oxygen consumption was measured from 3^rd^ instar larvae reared on control or 0.2 mg/ml CA supplemented food. Basal respiration values were normalized to the sample protein content prior calculating the change in respiration relative to control. Each group consisted of four to five replicates. ns, not significant; * p< 0.05; ** p< 0.01; *** p< 0.001.

**Figure S3.**
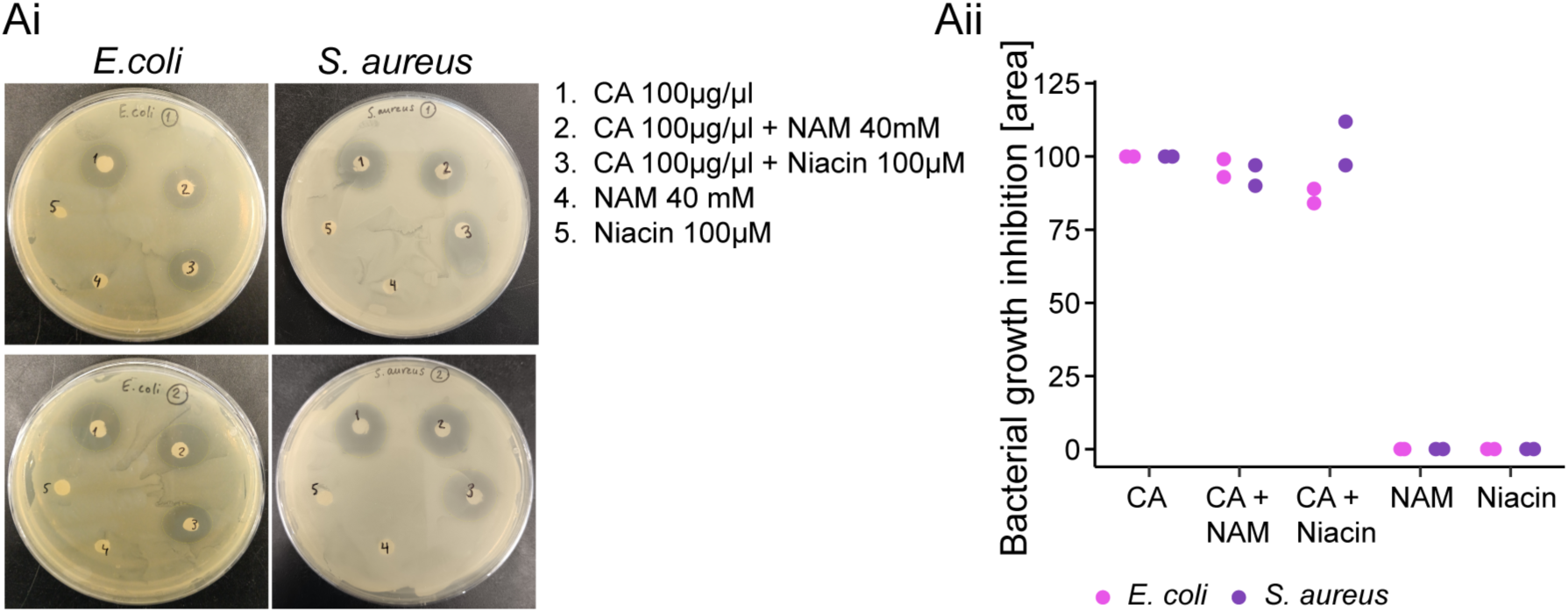
Antimicrobial activities of CA, NAM and niacin, individually and in combination. Antimicrobial activities of CA, CA + NAM, CA + niacin, NAM and niacin against Gram-negative bacterium *Escherichia coli* and Gram-positive bacterium *Staphylococcus aureus* based on the **Ai)** spot-on-lawn assay, and **Aii)** its quantification. Experiment was replicated on two plates shown in here.

**Figure S4.**
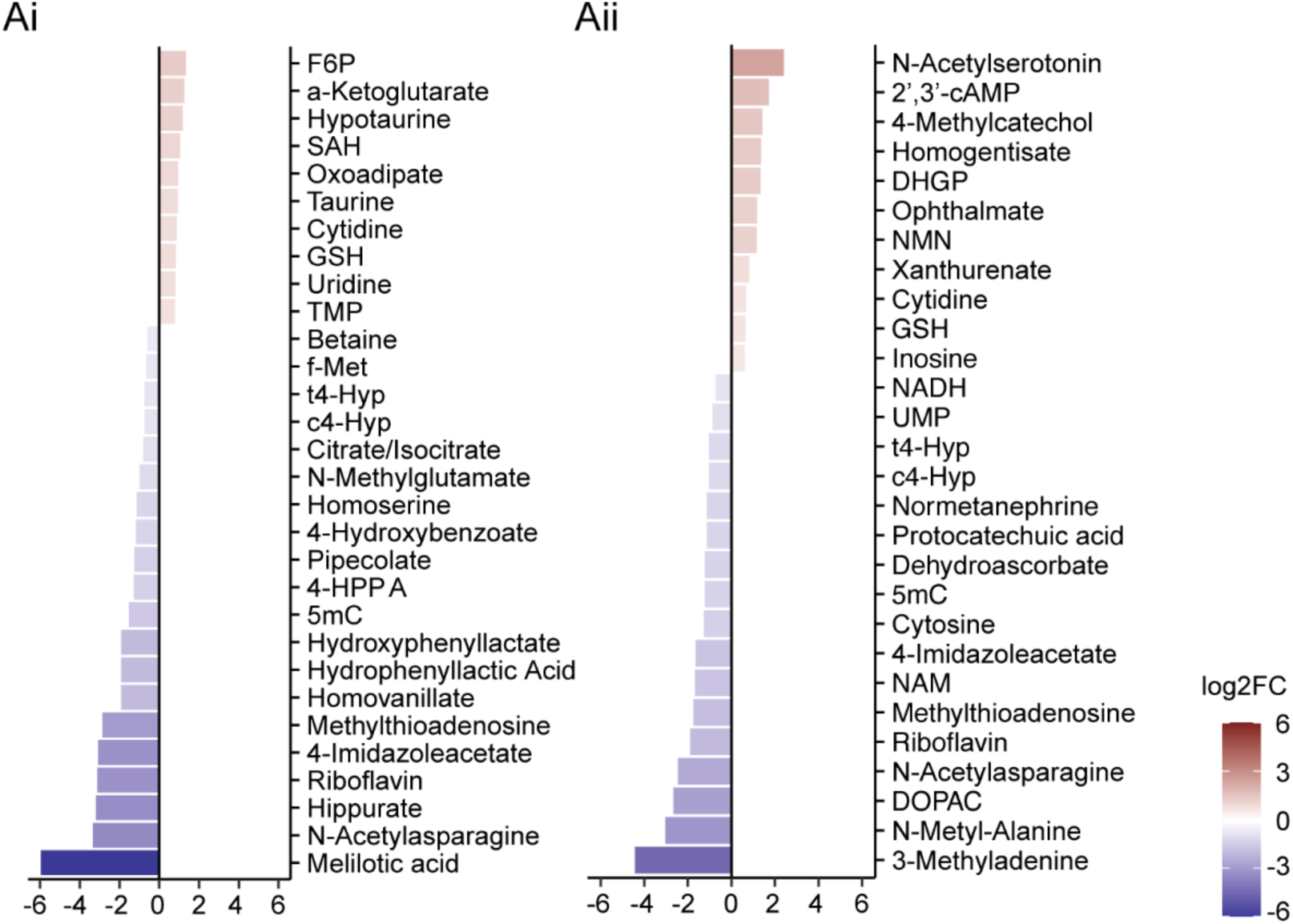
Effects of CA-feeding on ORT and nORT-mt765 larval metabolites. Most significantly differentially regulated metabolites in **Ai)** ORT (top 30) and **Aii)** nORT-mt765 (24) larvae fed with 2 mg/ml CA compared to their respective genetic controls fed on a control diet.

**Figure S5.**
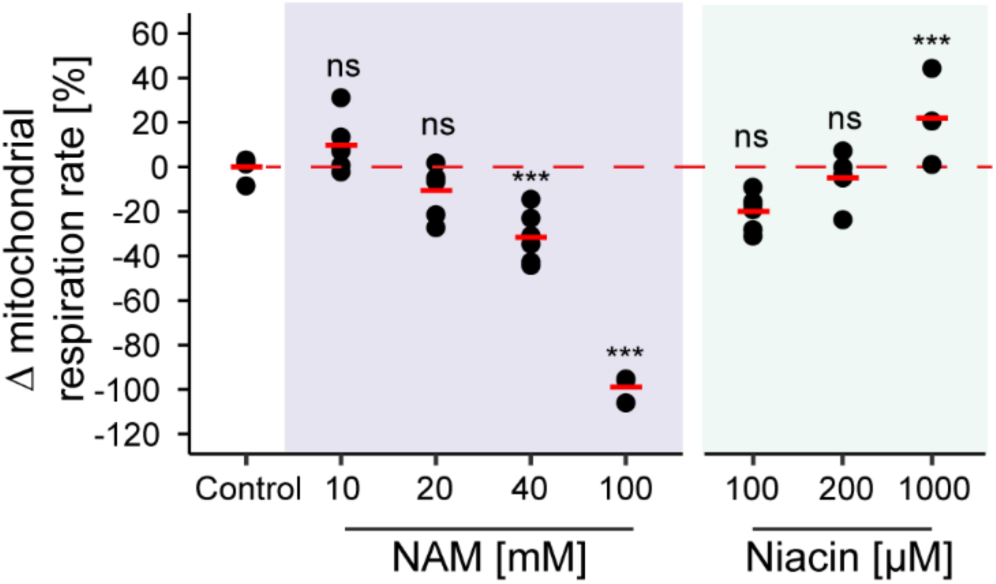
**Dose-specific effects of vitamin B_3_ vitamers on mitochondrial respiration**. HEK293T cells were exposed for 24 hours to different concentrations of nicotinamide (NAM; mM range) or niacin (µM range) and their effects on oxygen consumption were measured. Dashed red line refers to the average oxygen consumption level of the untreated HEK293T cells. ns, not significant; * p< 0.05; ** p< 0.01; *** p< 0.001.

